# Using experimental data and information criteria to guide model selection for reaction–diffusion problems in mathematical biology

**DOI:** 10.1101/444679

**Authors:** David J. Warne, Ruth E. Baker, Matthew J. Simpson

## Abstract

Reaction–diffusion models describing the movement, reproduction and death of individuals within a population are key mathematical modelling tools with widespread applications in mathematical biology. A diverse range of such continuum models have been applied in various biological contexts by choosing different flux and source terms in the reaction–diffusion framework. For example, to describe collective spreading of cell populations, the flux term may be chosen to reflect various movement mechanisms, such as random motion (diffusion), adhesion, haptotaxis, chemokinesis and chemotaxis. The choice of flux terms in specific applications, such as wound healing, is usually made heuristically, and rarely is it tested quantitatively against detailed cell density data. More generally, in mathematical biology, the questions of model validation and model selection have not received the same attention as the questions of model development and model analysis. Many studies do not consider model validation or model selection, and those that do often base the selection of the model on residual error criteria after model calibration is performed using nonlinear regression techniques. In this work, we present a model selection case study, in the context of cell invasion, with a very detailed experimental data set. Using Bayesian analysis and information criteria, we demonstrate that model selection and model validation should account for both residual errors and model complexity. These considerations are often overlooked in the mathematical biology literature. The results we present here provide a clear methodology that can be used to guide model selection across a range of applications. Furthermore, the case study we present provides a clear example where neglecting the role of model complexity can give rise to misleading outcomes.

## 1 Introduction

The development and testing of new theories to explain observations are keystones of the scientific method. In the biological sciences, mathematical models have become increasingly important to develop and test various hypotheses about putative mechanisms that drive biological processes. Furthermore, the interpretation of new biological data and the development of new experimental protocols is increasingly being enhanced through the use of mathematical models to explore questions of optimal data collection and optimal experimental design. However, for many applications in mathematical biology, there is a diverse range of valid modelling approaches that can be taken, and it is not always obvious which model is most appropriate for a particular application. Since all models are, by definition, a simplification of reality, the choice of modelling approach strongly depends on the purpose of the model and the set of modelling tools available. A complex model that accounts for many detailed mechanical, chemical and/or biological mechanisms may provide no additional insight than a simple phenomenological model in certain situations. Conversely, overly simple models that fail to capture key features of a particular process can lead to incorrect or incomplete conclusions.

An unresolved question in the field of mathematical biology is how to best determine when a model is sufficient. Informally, we might anticipate that the appropriate degree of model complexity will be related to the complexity and quality of the experimental data that we wish to model. However, judging the suitability of a model on its ability to match experimental data alone will always favour additional complexity over model simplicity. While this *best fit* approach is standard practice throughout the field of mathematical biology, we suggest that it is preferable instead to favour simple models unless additional complexity is warranted. In our work, we investigate and demonstrate techniques to select models, such as Bayesian analysis and information criteria, and give a practical illustration of the trade-off between consistency, fitness and complexity. We choose to focus on the question of selection between different continuum models that describe collective cell behaviour because this is a canonical modelling question of broad interest to the mathematical biology community.

In many biological applications, there can be multiple competing theories about the particular phenomenon that we might wish to study. In these situations, mathematical models can be used to objectively evaluate the validity of these potential theories by comparing model predictions with a set of experimental observations. Such an evaluation requires a robust, reproducible and objective framework for choosing the model, out of a set of candidates, that best explains the data. A typical approach to this model selection problem is to calibrate each candidate model to match a set of experimental data; the most traditional calibration approach being the maximum likelihood estimator (MLE) which is based on the minimisation of residuals. The model that fits the data “best” is selected as the model that best explains the observations. For example, Bianchi et al. (2016) use such an approach to evaluate several possible mechanisms that lead to wound healing failure through insufficient lymphangiogenesis. Mathematical models are also applied to interpret experimental data. For this purpose, model calibration is used to estimate model parameters that cannot be directly measured through experimentation (Sherratt and Murray, 1990). For example, estimates of the proliferation rate of cells can be obtained by calibrating a mathematical model to match data derived from *in vitro* cell culture assays (Jin et al., 2016b; Johnston et al., 2016; Maini et al., 2004). However, these parameter estimates depend implicitly upon the structure of the model that is used to interpret the experimental data.

### 1.1 Continuum models in cell biology applications

Continuum models of collective cell spreading and proliferation are used to study cancer, wound healing, embryonic development, and tissue engineering (Murray, 2002). However, the details of the models are diverse (Jin et al., 2016b; Simpson et al., 2011). The motility of cells can be modelled by: linear diffusion (Jackson et al., 2015; Maini et al., 2004; Sengers et al., 2007; Simpson et al., 2007; Swanson et al., 2003); nonlinear diffusion where the diffusivity increases with density (Bianchi et al., 2016; Flegg et al., 2009; Sengers et al., 2007), and nonlinear diffusion where the diffusivity decreases with density (Cai et al., 2007; King and McCabe, 2003); nonlinear advection to describe directed motility, such as chemotaxis (Bianchi et al., 2016; Flegg et al., 2010), haptotaxis (Marchant et al., 2001), and cell–cell adhesion (Armstrong et al., 2009). Similarly, the proliferation of cells is often modelled using a logistic source term (Maini et al., 2004; Sengers et al., 2007; Simpson et al., 2007), but there are also many other options for modelling carrying capacity-limited proliferation (Browning et al., 2017; Gerlee, 2013; Tsoularis and Wallace, 2002). Furthermore, the motility and proliferation of cells may also be coupled to diffusing chemical factors (Bianchi et al., 2016; Nardini et al., 2016; Savla et al., 2004; Sherratt and Murray, 1990). Despite this diverse range of modelling possibilities, few studies in the mathematical biology literature evaluate model uncertainty and many studies never consider any kind of model selection at all.

To illustrate these ideas we point to the work of Sherratt and Murray (1990) who present several continuum models of epidermal wound healing. To describe the motility of a cell population with density *C*, one model uses linear diffusion, with a constant nonlinear diffusivity function *D*(*C*) = *D*_0_, another model considers nonlinear diffusion so that the motility of cells depends on cell density, *D*(*C*) = *D*_0_*C^r^*, where *r* is an additional model parameter. Yet another model Sherratt and Murray (1990) consider includes linear diffusion with activator/inhibitor chemical regulation of cell proliferation. While Sherratt and Murray (1990) conclude that the chemical activator/inhibitor model provides the best match to the experimental data, they also conclude that nonlinear diffusion models with *r* = 1 or *r* = 4 also fit the data well, and even the simplest linear diffusion model, with *r* = 0, agrees with the data to some extent. Here, the model fit is directly proportional to the model complexity; hence the conclusions could be a result of overparameterisation (Akaike, 1974; Box, 1976). Therefore, a relevant question for us to address is: given complex biological data with many sources of uncertainty, how can we select models that provide a balance between model simplicity and agreement with data?

### 1.2 Challenges for model calibration

The study of the temporal growth, and the spatiotemporal spreading of cell populations provides us with a canonical area within the mathematical biology literature where there are many different types of continuum models available to interpret experimental data. For example, Sarapata and de Pillis (2014) catalog the most commonly used temporal growth models of tumors (Gerlee, 2013) and calibrate them against data from both *in vitro* and *in vivo* assays. Interestingly, Sarapata and de Pillis (2014) note that some of the MLE solutions lead to non-physical predictions about the carrying capacity density. Through application of a Bayesian approach, Warne et al. (2017) demonstrate that parameter uncertainty in such temporal models may be used to inform experimental design of proliferation assays. Sarapata and de Pillis (2014) and Warne et al. (2017) focus on temporal growth dynamics only, and they note that data obtained through standard experimental protocols may not contain sufficient information to resolve accurate and realistic estimates for all unknown parameters. Thus we expect that experimental design and model calibration requires detailed spatial data to compensate for increased model complexity when considering spatial models that describe the spatiotemporal spreading of cell populations. However, very few studies that calibrate spatial models of collective cell spreading use detailed cell density data. For example, Maini et al. (2004) and Sherratt and Murray (1990) simply use the location of the moving cell front to calibrate spatial reaction–diffusion models without considering detailed spatial cell density information. In both cases, Maini et al. (2004) and Sherratt and Murray (1990) use maximum likelihood estimation to calibrate these reaction–diffusion models. In these instances, the MLE provides no information on how identifiable the model parameters are from the moving front location alone. In a more recent study, Jin et al. (2016b) use detailed cell density profiles to identify parameter estimates based on the MLE solution for two commonly used models of collective cell spreading. The main point of Jin et al. (2016b) is to show that parameter estimates appear to depend strongly upon the initial density of cells in the experiments, and this result is at odds with our intuitive expectation because typical models implicitly assume that the parameters are independent of cell density. This indicates that a cautionary approach must be taken to reliably estimate parameters and compare models, and provide a partial explanation about why cell biology experiments are notoriously difficult to reproduce (Jin et al., 2016b).

The examples of Jin et al. (2016b) and Sarapata and de Pillis (2014) highlight a problem in model calibration and parameter estimation that is also discussed extensively by Slezak et al. (2010). That is, the common protocol in which candidate models are calibrated and compared using mathematical optimisation to determine the MLE from data can be misleading. Not only can this approach lead to biologically unrealistic parameter estimates or model behaviour (Jin et al., 2016b; Sarapata and de Pillis, 2014; Slezak et al., 2010) but comparison of maximum likelihood estimates is biased towards complex models that essentially overfit through an overabundance of free parameters (Box, 1976; Gelman et al., 2014; Johnson and Omland, 2004; Stoica and Selen, 2004). Furthermore, a traditional model selection process, based on maximum likelihood estimates, fails to capture uncertainty (Warne et al., 2017). There are four key sources of uncertainty when applying a mathematical model to interpret experimental data: 1) unknown model parameters that require statistical estimation; 2) uncertainty in the choice of model; 3) uncertainty that arises from stochastic fluctuations in the system dynamics; and 4) uncertainty resulting from systematic or measurement error in experimental work. Careful treatment of all of these sources of uncertainty is important to reliably validate theory and analyse data. Bayesian inference techniques are promising alternatives to MLE-based methods, since they can account for all relevant sources of uncertainty. Bayesian frameworks have been demonstrated to be highly effective at determining optimal experimental designs to minimise parameter uncertainty under the presence of systematic and measurement error (Browning et al., 2017; Johnston et al., 2016; Liepe et al., 2013; Parker et al., 2018; Vanlier et al., 2012; Warne et al., 2017).

The Bayesian view, however, has its challenges, such as, potential subjectivity of inference and lack of a clear decision process (Efron, 1986; Gelman, 2008a,b; Lambert, 2018). As a result, extensive statistical research to address these problems is an ongoing endeavour (Akaike, 1974; Gelman et al., 2014; Schwarz, 1978; Spiegelhalter et al., 2002), and there are now many alternative statistical techniques to traditional MLE-based methods or null hypothesis testing. Johnson and Omland (2004) review a number of common approaches in the context of ecological and evolutionary research, and Gelman et al. (2014) provide a more detailed review of model selection techniques from a statistical theory perspective.

### 1.3 Contribution

In this work we demonstrate how to apply a Bayesian framework to quantitatively compare and select a reaction– diffusion model of collective cell behaviour using detailed *in vitro* assay data. We show that the Bayesian view of datadriven model selection provides significantly more insight than traditional MLE approaches. A family of continuum reaction–diffusion models that describe cell motility and cell proliferation are evaluated from a Bayesian perspective through parameter uncertainty quantification. This exercise reveals important aspects of reliable model selection that are often not easily identified otherwise. We also evaluate a number of widely used decision-marking processes for model selection and compare the results with the intuition gained from the full Bayesian approach. In particular, we demonstrate, using detailed experimental data, that a trade-off between model fit, complexity and consistency can be obtained using a Bayesian framework. Furthermore, the techniques presented here valid for a wide class of models and are relevant for model comparison of in stochastic settings such as those considered by Johnston et al. (2016) and Matsiaka et al. (2018). Thus, we expect this work to be an exemplar of the Bayesian approach to the wider mathematical biology community.

## 2 Cell culture protocols

In cell biology, *in vitro* cell culture assays are commonly used to measure and observe the behaviour of cell populations in different environments. Typical examples are proliferation assays (Browning et al., 2017), scratch assays (Jin et al., 2016b) and invasion assays (Haridas, 2017). In this work, we will focus on the scratch assay (Liang et al., 2007), however, it is important to note that our methods are widely applicable to other assay types. Jin et al. (2016b) provide a particularly detailed scratch assay data set using the PC-3 prostate cancer cell line. Each well in a 96-well tissue culture plate are identically populated with a particular initial number of PC-3 cells. Cells are left to attach to the substrate for a small amount of time so that a series of uniform monolayers have formed. Then, identical scratches, approximately 0.5 mm in width, are made in the monolayers. Images of the scratched region are then captured at regular time intervals for a total duration of 48 hours. A particularly insightful feature of the protocol used by Jin et al. (2016b) is that they performed multiple experiments to demonstrate how variation in the initial density of cells in the monolayer affect cell invasion. Experiments were perform by initially placing either 10, 000, 12, 000, 14, 000, 16, 000, 18, 000 or 20, 000 cells into the wells of the 96-well plate. For brevity, in our study, we focus on data from the experiments initialised with 12, 000, 16, 000 and 20, 000 cells per well. Some example images from these datasets are summarised in Figure 1.

**Fig. 1.**
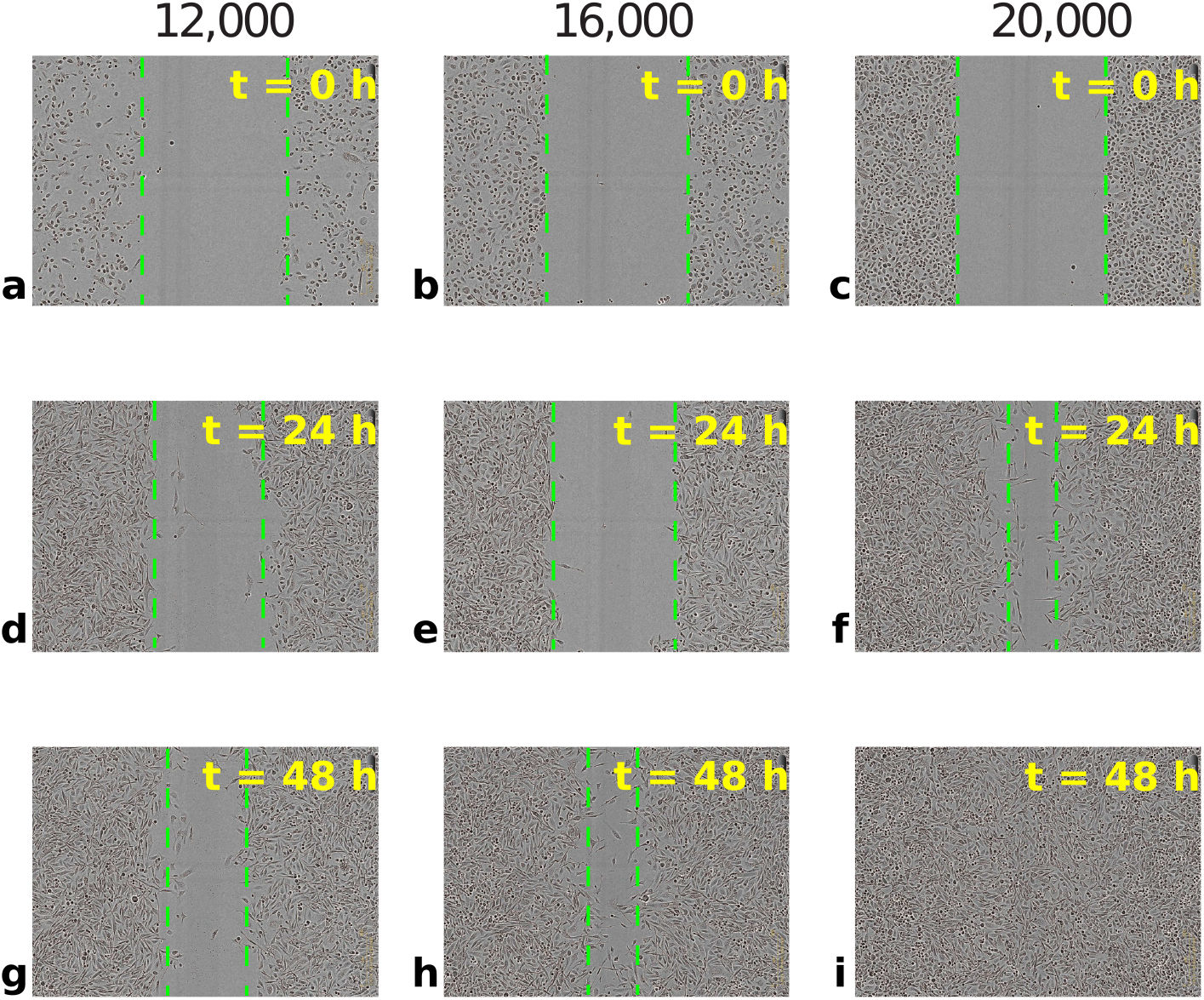
Example scratch assay data using the PC-3 prostate cancer cell line. Each column presents one experiment using an initial population of: (a),(d) and (g) 12, 000 cells; (b),(e) and (g) 16, 000 cells; (c),(f) and (i) and 20, 000 cells. The dimensions of each image are 1.43 *×* 1.97 [mm]. Images are shown at 24 [h] time intervals, however, the data is captured at 12 [h] intervals. Images are reproduced from Jin et al. (2016b), with permission.

The data in Figure 1 demonstrate that the scratch closure rate depends on the initial density of cells. For the low density initial condition of 12,000 cells per well, the scratch remains more than half its original size after 48 hours (Figure 1(a),(d) and (g)). In contrast, for the medium initial density of 16,000 cells per well, the scratch area is noticeably smaller at 48 hours, but it is still not closed (Figure 1(b),(e) and (g)). Only the high initial density initial condition of 20,000 cells per well leads to complete scratch closure after 48 hours (Figure 1(c),(f) and (i)). Most scratch assay protocols do not consider varying the initial density of cells, and those studies that use mathematical models to interpret experimental results from a scratch assay focus only on temporal data describing the position of the leading cell front (Sherratt and Murray, 1990; Maini et al., 2004), indicated by the dashed green line in Figure 1(a)–(h). The study of Jin et al. (2016b) is unique since they provide detailed cell density profiles from multiple experimental replicates for each initial condition considered. These high resolution data enable us to pose and explore new questions about the applicability of some commonly used continuum reaction–diffusion models to describe this dataset. We utilise a subset of the original PC-3 scratch assay data (See Jin et al. (2016b)) to continue this line of reasoning using Bayesian analysis.

## 3 Continuum models of cell motility and proliferation

Fundamental features associated with cell invasion processes are the motility and proliferation of cells. Many different intracellular and intercellular mechanisms are relevant to both motility and proliferation, depending on the specific biological process. In the biological literature, it is not always clear which mechanisms are most important or relevant in a particular situation. Furthermore, it may be difficult to identify the most appropriate mathematical model, especially when a variety of models fit the experimental data both qualitatively and quantitatively. This is particularly true in the area of modelling of epidermal wound healing (Maini et al., 2004; Murray, 2002; Sherratt and Murray, 1990; Simpson et al., 2011).

### 3.1 Modelling cell populations with reaction–diffusion equations

Continuum models are routinely used to describe the evolution of a population of cells that undergo collective cell spreading and proliferation. Such models are often based on partial differential equations (PDEs) that are reaction–diffusion equations of the form

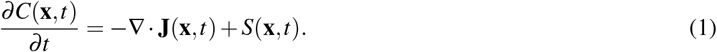

Here, *C*(**x***, t*), is the cell density at position, **x**, and time *t*, **J**(**x**, **t**) is the cell population density flux vector and *S*(**x***, t*) is a reaction term representing cell proliferation and loss. Both **J**(**x***, t*) and *S*(**x***, t*) are often functions of the cell density, *C*(**x***, t*), the cell density gradient vector, ∇*C*(**x***, t*), or both; there is also usually a dependence on model parameters that must be calibrated using experimental data. The functional forms of **J**(**x***, t*) and *S*(**x***, t*) used for modelling vary across diverse applications in tumor growth/invasion, wound healing, and embryology. However, there are some common choices. For the growth function, *S*(**x***, t*), Gerlee (2013) and Sarapata and de Pillis (2014) describe many of the key growth models in the context of tumor growth. A variety of options for the flux, **J**(**x***, t*), are discussed by Murray (2002), and Simpson et al. (2006) perform a simulations of these options and compare qualitatively with experimental data.

Of the available growth models, the most fundamental is the logistic growth model,

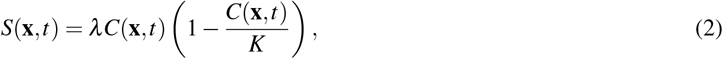

where *λ >* 0 is the rate of cell proliferation, *K >* 0 is the carrying capacity density, that is, the cell population density at which contact inhibition reduces the net population growth to zero. The logistic growth model is frequently used to describe cell growth as it is the most fundamental model that describes the effect of contact inhibition of proliferation (Warne et al., 2017), however, general forms can also be considered in cell biology applications (Browning et al., 2017; Sarapata and de Pillis, 2014; Tsoularis and Wallace, 2002). For the flux, **J**(**x***, t*), the most common choice is Fickian diffusion (Maini et al., 2004; Sherratt and Murray, 1990),

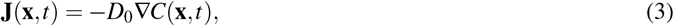

where *D*_0_ *>* 0 is a constant cell diffusivity. This formulation of the flux models cells for which motility is not affected by cell density, that is, cells are behaving like Brownian particles.

When logistic growth (Equation (2)) and Fickian diffusion (Equation (3)) are substituted into (Equation (1)) we obtain

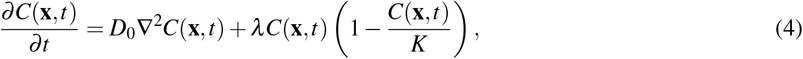

which, in one dimension, is known as the Fisher–Kolmogorov–Petrovsky–Piscounov model (Fisher–KPP) (Murray, 2002). The Fisher–KPP model has been applied in many biological contexts (Murray, 2002). In cell biology, common applications include the modelling of wound healing (Sherratt and Murray, 1990), tissue engineering (Sengers et al., 2007), tumor growth (Swanson et al., 2002), cancer treatment (Jackson et al., 2015), and embryonic development (Simpson et al., 2007).

Maini et al. (2004) demonstrate there is experimental evidence for the standard Fisher–KPP model in the dynamics of human peritoneal mesothelial cells. However, the standard Fisher–KPP model is often modified to capture application specific features; most changes relate to the form of *S*(**x***, t*) and **J**(**x***, t*) in Equation (1). Savla et al. (2004) modify the proliferation function to account for stretching that occurs in bronchial epithelial cells. The invasion of embryonic neural crest cells through the intestine is modelled by Simpson et al. (2007) by considering a multi-species version of Equation (4) where both species contribute to the carrying capacity. Swanson et al. (2003) model the growth and invasion of gliomas using a spatially heterogeneous diffusivity to account for grey matter and white matter regions of the brain. Inclusion of growth factors and chemical gradients is also often handled through coupling activator/inhibitor dynamics to models of chemical diffusion (Nardini et al., 2016; Sherratt and Murray, 1990).

A very common modification to the standard Fisher–KPP model is to incorporate density dependent diffusion of the form **J**(**x***, t*) = *−D*(*C*(**x***, t*))∇*C*(**x***, t*), where *D*(*C*(**x***, t*)) is the nonlinear diffusivity function. Often, *D*(*C*(**x***, t*)) is chosen to be a monotonically increasing function with *D*(0) = 0 (Flegg et al., 2010; Simpson et al., 2011; Sengers et al., 2007). It is also intriguing that other studies choose to focus on monotonically decreasing nonlinear diffusivity functions (Cai et al., 2007). If *D*(*C*(**x***, t*)) = *D*_0_*C*(**x***, t*)*/K*, where *D*_0_ is the cell diffusivity, we obtain the Porous Fisher model (Gurney and Nisbet, 1975; Murray, 2002):

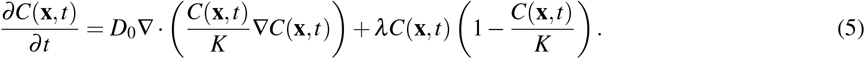

Unlike the Fisher–KPP model (Equation (4)), diffusion is not solely derived by the random movement of cells. Instead, cells exhibit movements that are directed away from crowded areas (Gurney and Nisbet, 1975), with a direct linear relationship between motility and density. Some studies have also considered *D*(*C*(**x***, t*)) = *D*_0_(*C*(**x***, t*)*/K*)^*r*^, which can be thought of as a Generalised Porous Fisher equation (Sherratt and Murray, 1990; Witelski, 1995),

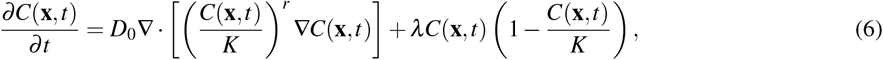

where *r* is a constant that controls the density avoidance/attraction behaviour of cells. Here, the Fisher–KPP and Porous Fisher models are recovered with *r* = 0 and *r* = 1, respectively. In applications, *r* is often selected quite arbitrarily (Jin et al., 2016b; Sherratt and Murray, 1990) and there is little theory enabling its biological interpretation (Simpson et al., 2011).

The interpretation of the power, *r*, indeed deserves further discussion. In effect, it models a nonlinear relationship between the motility of cells and the cell density (Simpson et al., 2011). For *r >* 1, the relationship is superlinear; the cell motility slowly increases with cell density at lower densities (Sherratt and Murray, 1990), but then rapidly increases at higher densities. For 0 *< r <* 1, we have a sublinear relationship, that is, cells increase in motility faster at low densities (Jin et al., 2016b). Some have also considered the case of “fast diffusion” (*r <* 0) where cells become increasingly motile as the density decreases (King and McCabe, 2003).

For certain special choices of boundary conditions and initial conditions, the Fisher–KPP, Porous Fisher and Generalised Porous Fisher models are known to have travelling wave solutions. These solutions are of general mathematical interest and have been extensively studied (Harris, 2004; Witelski, 1995). However, since travelling waves only occur as in the long-time limit and require rather special initial conditions, travelling waves are rarely observed experimentally (Jin et al., 2016b; Vittadello et al., 2018). Therefore, we do not consider connecting any kind of travelling wave solutions with experimental data in this work.

Different values of *r* also result in qualitatively different wave fronts (Murray, 2002). Figure 2(c), (g), (k) and (o) compares the evolution of the Generalised Porous Fisher model in one spatial dimension for initial and boundary conditions known to lead to travelling waves; parameters are selected to correspond to similar wave speeds. The travelling wave of the Fisher–KPP model (Figure 2(c)) has no distinct leading edge, since *C*(*x, t*) *>* 0 for all *x*. However, the Porous Fisher model (Figure 2(k)) exhibits a distinct interface, sometimes called the contact point, separating regions of zero and non-zero density. This is the case for any *r >* 0. The shape of the wave front also changes with *r*. In particular, the wave front is concave upward in Figure 2(a)–(h) and concave downward in Figure 2(i)–(p). This concavity change can be analysed using the exact solution to the Generalised Porous Fisher model (Equation (6)) when *λ* = 0 and the initial density is a Dirac delta function. This exact solution (Figure 2(a), (e), (i) and (m)) can be used to show that the wave front is concave upward for *r <* 1 and concave downward for *r ≥* 1 (see Appendix A and Murray (2002)). Aside from this exact solution (Figure 2(a), (e), (i) and (m)), all solutions in Figure 2 are computed numerically (see Appendix B for details on the numerical scheme that includes the spatial and temporal step sizes used throughout this work).

**Fig. 2.**
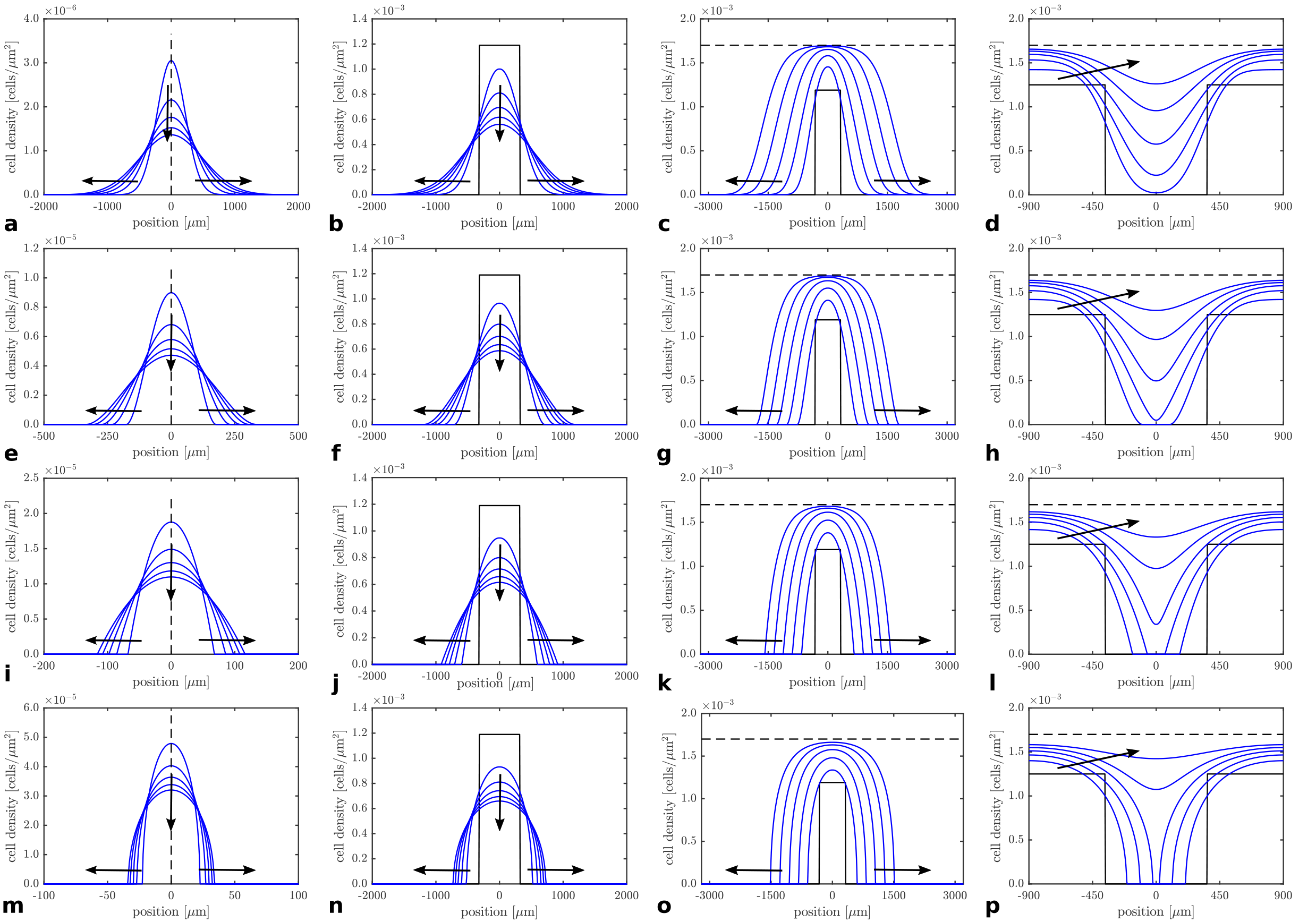
Evolution of the solution of the Generalised Porous Fisher equation (solid blue) for various parameter values and initial conditions. The various initial conditions are shown (solid black). Arrows indicate the direction of increasing *t* and profiles are shown at regular time intervals. Each row highlights the role of the exponent *r*: (a)–(d) *r* = 0; (e)–(h) *r* = 1*/*2; (i)–(l) *r* = 1; and (m)–(p) *r* = 2. Each column corresponds to a different initial conditions: (a), (e), (i) and (m) show the exact solution to the diffusion only problem with a delta function initial condition and *λ* = 0 shown at 24-hour time intervals; (b), (f), (j) and (n) show the solution of the diffusion-only problem for a spatially-extended initial condition, *λ* = 0 shown at 24-hour time intervals; (c), (g), (k) and (o) show the same initial condition and time intervals, but with logistic proliferation with proliferation rate *λ >* 0 and carrying capacity density *K* (dashed black); (d), (h), (l) and (p) show a simplified, but typical, wound healing configuration shown at 12-hour time intervals.

As stated previously, formal travelling wave solutions are almost never observed experimentally. Typical scratch assays initial configurations lead to two opposingly-directed fronts, such as the profiles in Figure 2(d), (h), (l) and (p) showing the evolution of the Generalised Porous Fisher model for several values of *r*. Note that, for typical scratch widths, wound closure takes place before travelling waves have an opportunity to form (Jin et al., 2016b; Vittadello et al., 2018). However, the value of *r* still impacts the wound closure rate and the shape of the moving font.

### 3.2 Model comparison

Traditional approaches to model calibration are usually based on regression (Johnson and Omland, 2004). Suppose we have data that consists of *n* experimental observations, 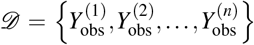 and a mathematical model, *y* = *M*(*x*; ***θ***), parameterised by ***θ*** *∈* ***Θ***, where ***Θ*** is a *k*-dimensional space of valid parameter combinations. The model makes predictions, 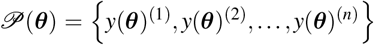, with *y*(***θ***)^(*i*)^ = *M*(*x*^(*i*)^; ***θ***) for *i* = 1, 2*,…, n* where *x*^(1)^*, x*^(2)^*,…, x*^(*n*)^ are model inputs; for example, a model input may include initial conditions, boundary conditions, or the spatiotemporal position of model predictions. The regression parameters, ***θ̂***, are obtained through minimisation of the residual error, that is,

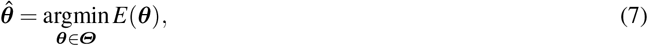

where *E*(***θ***) is the residual error,

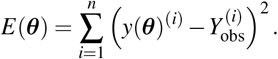

In Equation (7), ***θ̂*** corresponds to the MLE under the assumption of Gaussian observational error. Non-linear mathematical optimisation techniques are applied to obtain numerical solutions to Equation (7). One of the limitations of the MLE is that only a point estimate of the parameters is obtained, although bootstrapping may be applied to obtain confidence intervals (Gelman et al., 2004). While this can be sufficient, without more effective handling of uncertainty in modelling assumptions or experimental setup, the estimate can be biologically unrealistic (Slezak et al., 2010).

Based on detailed density data from a scratch assay (Figure 3a), it is unclear whether linear or nonlinear diffusion is most relevant. Sherratt and Murray (Sherratt and Murray, 1990) find that linear Fickian diffusion (*r* = 0) with chemically regulated proliferation provided a lower residual error than non-linear diffusion (with *r* = 4) using mammalian epidermal wound closure data. However, Sherratt and Murray (1990) never considered varying the initial cell density in the experiments or model simulations. Through multiple model calibrations using data with a range of initial densities, Jin et al. (2016b) demonstrates that estimates of *D*_0_ are not constant under changes in initial cell density, suggesting that the diffusion of PC-3 prostate cancer cells is density dependent.

**Fig. 3.**
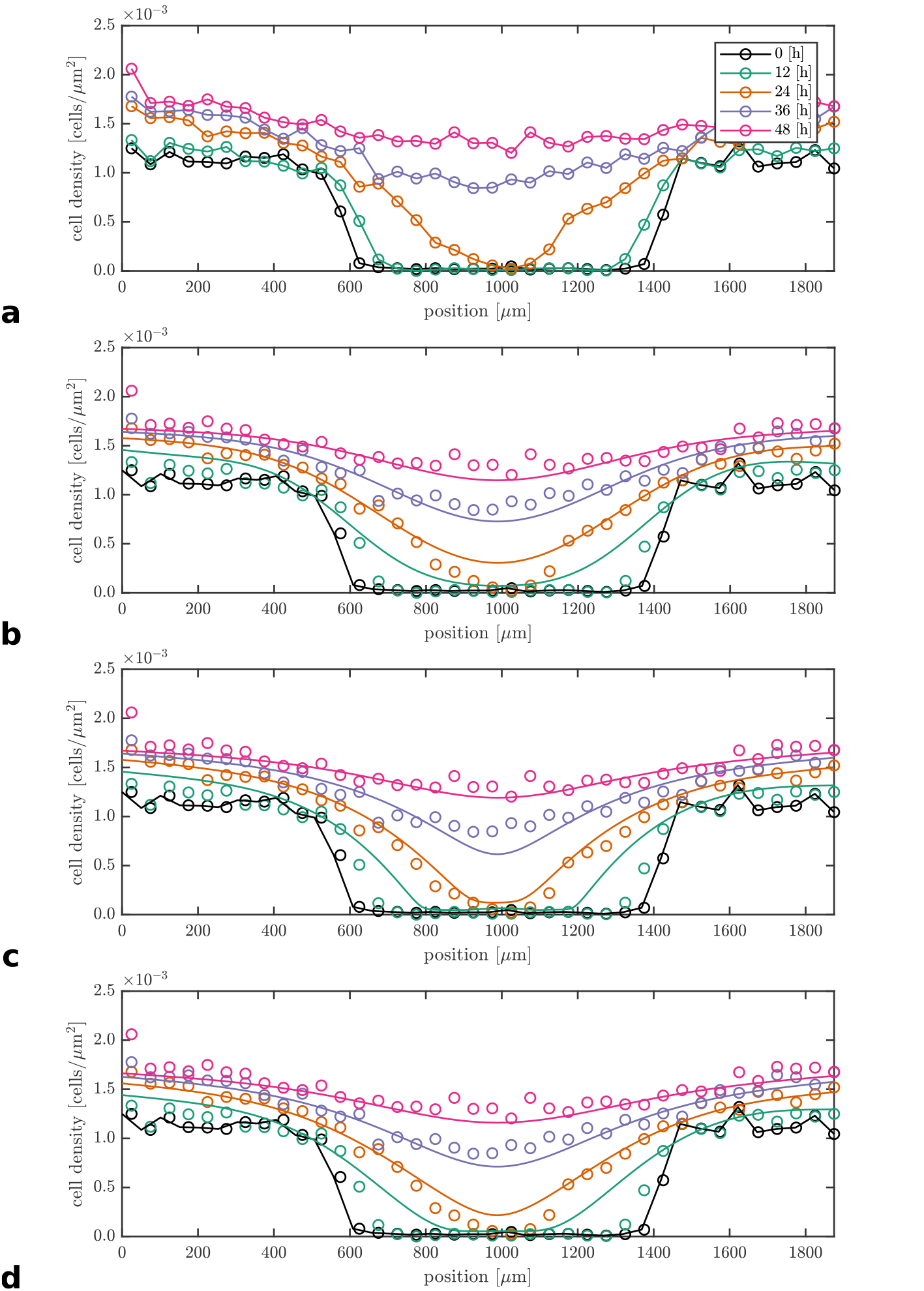
Calibration of the (b) Fisher–KPP, (c) Porous Fisher, and (d) Generalised Porous Fisher models against (a) PC-3 scratch assay data seeded with 20, 000 cells (as obtained by Jin et al. (2016b)). (a)–(d) The experimental data are shown at times *t* = 0 [h] (black circles), *t* = 12 [h] (green circles), *t* = 24 [h] (orange circles), *t* = 36 [h] (light blue circles), and *t* = 48 [h] (magenta circles); (a) linear interpolation of data are also shown. The obtained MLE parameter estimates are: (b) *D*_0_ = 1030 [*μ*m^2^/h], *λ* = 6.4 *×* 10^*−*2^ [1/h] for the Fisher–KPP model; (c) *D*_0_ = 2900 [*μ*m^2^/h], *λ* = 6.4 *×* 10^*−*2^ [1/h] for the Porous Fisher model; and (d) *D*_0_ = 2160 [*μ*m^2^/h], *λ* = 5.8 *×* 10^*−*2^ [1/h], *r* = 5.2 *×* 10^*−*1^ for the Generalised Porous Fisher model. The carrying capacity density is *K* = 1.7 *×* 10^*−*3^ [cells/*μ*m^2^], determined using cell densities at *t* = 48 [h] within 200 [*μ*m] from the left and right boundaries. The initial density profile (solid black) is determined through linear interpolation of the data at time *t* = 0 [h].

The work of Jin et al. (2016b) highlights the need to calibrate models over multiple datasets to effectively compare them. Figure 3(b) and (c) show scratch assay density profiles measured at 12-hour intervals superimposed on plots of the solutions of the Fisher–KPP model and the Porous Fisher model using the parameter estimates reported by Jin et al. (2016b). They used mathematical optimisation (Jin et al., 2016b) to find the MLE for the parameters, ***θ*** = {*D*_0_*, λ*}, assuming *K* is fixed at a value that they estimate independently using only regions far from the scratch area at late time, *t* = 48 [h], so that the packing density they observed was close to the maximum possible packing density. Results in Figure 3 show that both models fit the data well. The minimised residual error, *E*(***θ****̂*), respectively, is *E*(***θ̂***) = 2.48 *×* 10^*−*6^ for the Fisher–KPP model, and *E*(***θ̂***) = 2.58 *×* 10^*−*6^ for the Porous Fisher model. If model selection were to be based on the minimum residual error, the conclusion would be that the Fisher–KPP model explains the data the best, even if by a small margin. However, we will show that this is an overly simplistic conclusion in this case. Since the Fisher–KPP model (Equation (4)) and the Porous Fisher model (Equation (5)) are both special cases of the Generalised Porous Fisher model (Equation (6)), the Generalised Porous Fisher model cannot have a higher residual error than either of the two special cases. For example, the calibrated Generalised Porous Fisher model, shown in Figure 3(d), has a residual error of *E*(***θ̂***) = 2.47 *×* 10^*−*6^. However, we demonstrate that decreased residual error comes at a cost that is totally obscured by taking this standard approach to model selection. As we show, the trade off between model fitness and model complexity can be made very clear using Bayesian techniques.

It should be noted that there are other mechanisms that could be added to enhance the data fit. For example, diffusion of growth factors and/or chemotaxis mechanisms could be included in the suite of potential models that we apply to the experimental data (Bianchi et al., 2016; Nardini et al., 2016; Sherratt and Murray, 1990). Just as with the Generalised Porous Fisher model, this always leads to extra parameters that will enable the model to fit the data better. We suggest this improvement is meaningless without carefully considering the uncertainty in the parameter estimates and the increased model complexity, and we will provide further discussion on this point in the Conclusions section.

## 4 A Bayesian framework for model comparison

In this section, we analyse the PC-3 scratch assay density profiles from a Bayesian perspective. We demonstrate that, despite the MLE approach giving preference to the standard Fisher–KPP model, there are other reasons to consider the Porous Fisher model as preferable in this case. This demonstration indicates that it is of benefit to include Bayesian uncertainty quantification as a standard technique for model calibration and validation in biological applications.

### 4.1 Fundamentals of Bayesian analysis

The Bayesian approach is to consider unknown model parameters as random variables with their respective probability distributions representing what is known about the parameters (Efron, 1986; Gelman et al., 2014; Lambert et al., 2018). The conditional probabilities of the parameters given experimental observations represent the new knowledge obtained from an experiment under the assumption of a given model.

Mathematically, this is expressed though Bayes’ Theorem (Gelman et al., 2014),

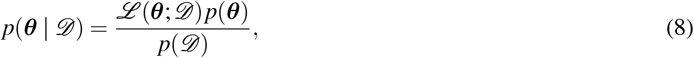

where ***θ*** is the vector of unknown parameters that exist in some parameter space, ***Θ***, and *𝒟* is the set of observations within some space of possible outcomes, *𝔻*. The *prior* probability density, *p*(***θ***), represents any *a priori* knowledge preceding observations, the likelihood, *ℒ* (***θ***; *𝒟*), is the probability density of the observations, *𝒟*, and *p*(***θ*** *| 𝒟*) is the resulting *posterior* probability density representing new knowledge of the parameters after including observations. The *evidence*, *p*(*𝒟*), is a probability density function (PDF) for the observations over all parameters, that is, 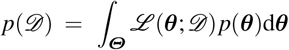; from a practical perspective, the evidence is a normalisation constant.

Conceptually, the prior and the likelihood encode assumptions; the former is related to assumed knowledge of parameters and the latter to the underlying mathematical model. One criticism of the Bayesian approach is that the requirement of a prior leads to subjectivity since a strict “zero-information” test cannot be formally defined (Efron, 1986). On the other hand, the Bayesian approach is capable of dealing with arbitrarily complex models and priors, thus providing a very general and consistent analysis framework (Efron, 1986).

The Bayesian posterior PDF provides a natural way to describe uncertainty in parameter estimates. In this context, the uncertainty in the *i*th parameter estimate, ***θ***_*i*_, is defined as the variance of ***θ***_*i*_ with respect to the posterior distribution, where

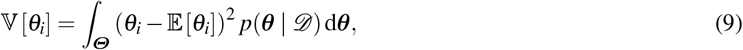

and

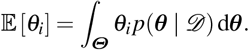

Bayesian methods have been shown to be highly effective at informing experimental design and parameter inference (Browning et al., 2017; Johnston et al., 2016; Lambert et al., 2018; Silk et al., 2014; Vanlier et al., 2012; Warne et al., 2017). Through the Bayesian formulation, experiments can be designed so as to minimise the level of uncertainty in estimates of a given parameter set. Furthermore, routine experimental protocols can be analysed to identify areas for potential improvement (Warne et al., 2017).

### 4.2 Scratch assay data informs continuum model comparison

To interpret scratch assay data we let *C*_obs_(*x, t*) be the observed cell density at position *x* (in the one-dimensional profile) and time *t*. We assume that observations are subject to additive noise,

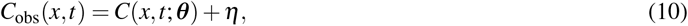

where *C*(*x, t*; ***θ***) is the true density as determined through the assumed continuum model with parameters, ***θ***, as discussed in Section 3, and *η* represents the combination of measurement error, systematic error and stochastic fluctuations. For simplicity, we treat the error, *η*, as Gaussian noise with mean zero, and known variance *σ*^2^, that is, *η* ∼ *𝒩* (0*, σ*^2^). Furthermore, such an assumption ensures that our likelihood formulation corresponds to the likelihood implied by the MLE interpretation of the non-linear regression approach to model calibration (Equation (7)). However, it should be noted that the Bayesian techniques we apply here do not require this assumption, nor is it a requirement that *σ* be known.

We also specify, for ease of description, that *C*_obs_(*x*, 0) = *C*(*x*, 0; ***θ***), that is, perfect observation is possible for the initial condition. This is a reasonable and realistic assumption to make. However, it should be noted that our framework can be extended to deal with cases of observation error in the initial conditions. The treatment of noise in the initial condition observation error is an interesting point for discussion, so we provide further results and details in Appendix D. Importantly, parameter uncertainty is amplified in this case. See also Jin et al. (2016b) and Warne et al. (2017) for further details.

Processed scratch assay data are of the form, 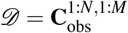, where 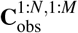 is an *N × M* matrix with elements that are observations, as given by Equation (10), at *NM* position-time pairs taken from the Cartesian product of *N* spatial points *x*_1_*, x*_2_*,…, x_N_* with *M* temporal points *t*_1_*, t*_2_*,…, t_M_*. That is,

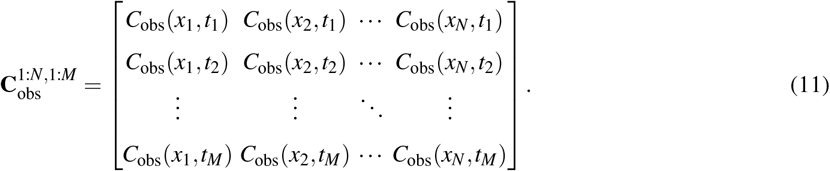

The PC-3 cell line scratch assay datasets, described in Section 2, are derived from microscopy images that are taken at times *t*_1_ = 0 [h], *t*_2_ = 12 [h], *t*_3_ = 24 [h], *t*_4_ = 36 [h], and *t*_5_ = 48 [h]. Densities are computed, for each point in time, using rectangular areas with centerlines at *x*_1_ = 25*, x*_2_ = 75*, x*_3_ = 125*,…, x*_38_ = 1925 [*μ*m]. That is, *N* = 38 and *M* = 5. These derived data are provided by Jin et al. (2016b), and are more detailed that the data used by Maini et al. (2004) and Sherratt and Murray (1990). However, the data used by Maini et al. (2004) and Sherratt and Murray (1990) are still more detailed than many studies in which microscopy images alone, without any quantitative measurements of density or spatial position, are used to illustrate the outcomes of a scratch assay.

First, we consider the Fisher–KPP (Equation (4)) and Porous Fisher models (Equation (5)). The task is to construct a Bayesian posterior PDF for the parameters ***θ*** = [*D*_0_*, λ, K*] given each of the initial conditions and under the assumption of each model. The resulting posterior PDFs may be compared visually to get intuition on the appropriateness of each model given the PC-3 data. Most analyses of cell motility and proliferation assume the carrying capacity density *K* is a known parameter (Warne et al., 2017), however, this is usually an approximation that is required due to short assay timescales (Warne et al., 2017). In the case of the data of Jin et al. (2016b), *K* is more appropriately considered as unknown since the data captures more of the long-time effects of contact inhibition (Sarapata and de Pillis, 2014; Warne et al., 2017).

To keep subjective bias to a minimum, the aim is to assume as little as possible within the prior distributions, that is we wish them to be uninformative. To this end, we select uniform prior distributions for each of the three parameters such that the support extends well beyond biologically viable ranges. In the literature, typical ranges for each parameter are: *D*_0_ = 155–6500[*μ*m^2^*/*h]; *λ* = 0.01–0.07[1/h]; and *K* = 1.5 *×* 10^*−*3^–2.0 *×* 10^*−*3^[cells*/μ*m^2^] (Browning et al., 2017; Maini et al., 2004; Jin et al., 2016b). Therefore, we assume as priors *D*_0_ *∼ 𝒰* (0*, D*_max_), *λ ∼ 𝒰* (0*, λ*_max_), and *K ∼ 𝒰* (0*, K*_max_) where *D*_max_ = 10^5^ [*μ*m^2^*/*h], *λ*_max_ = 1 [1*/*h] and *K*_max_ = 7 *×* 10^*−*3^ [cells*/μ*m^2^]. As a joint PDF, we have

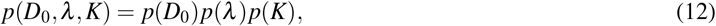

where

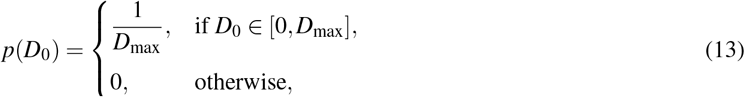

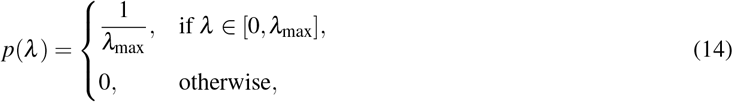

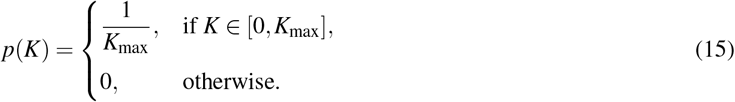

It is important to note, however, that uniform priors are not always uninformative and care must be taken (Efron, 1986).

Under the aforementioned assumption of independent Gaussian observation error on the data (Equation (10) and Equation (11)), the likelihood function is

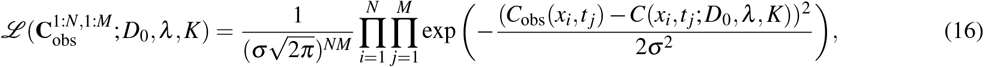

where *C*(*x_i_, t _j_*; *D*_0_*, λ, K*) is the solution of the continuum model of interest, computed numerically (see Appendix B) at point *x_i_* and time *t _j_*, given values for *D*_0_, *λ* and *K*. The model will be either the Fisher–KPP model (Equation (4)) or Porous Fisher model (Equation (5)). The observation error is such that *σ ≈* 10^*−*5^ (cells/*μ*m^2^), as obtained from previous studies (Jin et al., 2016b; Warne et al., 2017). Comparing the non-linear regression problem in Equation (7) with this likelihood (Equation (16)) reveals that the MLE parameter set in Equation (7) corresponds to the mode of the likelihood as a function of the parameter vector, ***θ*** = [*D*_0_*, λ, K*].

Substitution of the prior PDF (Equation (12)) and the likelihood (Equation (16)) into the right-hand side of Bayes’ Theorem (Equation (8)) yields the posterior PDF,

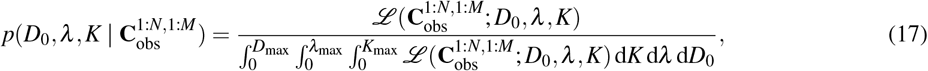

when [*D*_0_*, λ, K*] ∈ [0*, D*_max_] *×* [0*, λ*_max_] *×* [0*, K*_max_], otherwise 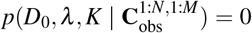. From Equation (17), we can construct the posterior marginal PDFs,

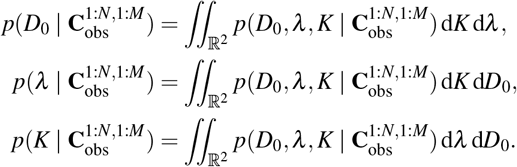

The posterior marginal PDFs represent the uncertainty in a single parameter taken over all possibilities of the remaining parameters. The integrals are best computed using Monte Carlo integration. Specifically, we apply approximate Bayesian computation (ABC) rejection sampling (Sunnåker et al., 2013) to obtain samples from the joint posterior density, then apply Monte Carlo integration with kernel smoothing to estimate the posterior marginal densities (Silverman, 1986). We leave the computational details to Appendix C, however, it is important to note that, for deterministic models with additive noise, the ABC rejection sampler can be shown to be a exact (Wilkinson, 2013).

We compare the posterior marginal PDFs obtained under the assumption of the Fisher–KPP model (Equation (4)), conditioned on data as given in Equation (11), against those obtained under the assumption of the Porous Fisher model (Equation (5)). Posterior marginal PDFs are computed using ABC rejection sampling to generate *n* = 50, 000 samples from the joint posterior distribution as described in Appendix C. The results are presented in Figure 4. Leaving more quantitative analysis for Section 5, we discuss here the qualitative aspects that are essential for, not only model selection and comparison, but also experimental design.

**Fig. 4.**
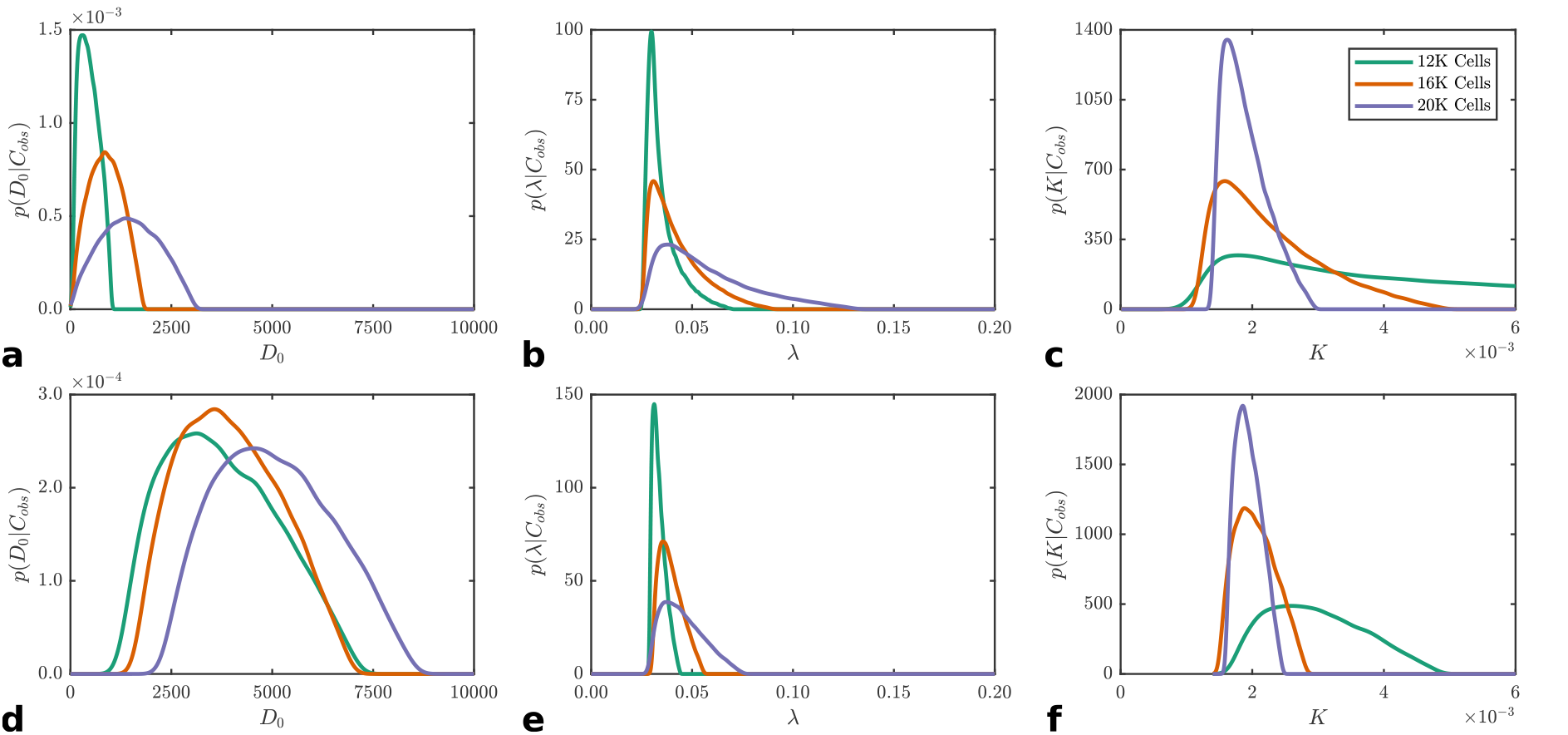
Marginal posterior probability densities obtained through Bayesian inference on the PC-3 scratch assay data under the (a)–(c) Fisher– KPP model and (d)–(f) Porous Fisher model. The spread in marginal densities demonstrate the degree of uncertainty in the diffusivity, *D*_0_, the proliferation rate, *λ*, and the carrying capacity, *K*, for each model under the three different initial conditions; 12, 000 cells (solid green), 16, 000 cells (solid orange), 20, 000 cells (solid purple).

The first point of interest is the trade-off between uncertainty in the proliferation rate, *λ*, and the carrying capacity density, *K*. The uncertainty in *λ* increases as the initial cell density increases (Figure 4(b) and (e)), however, the reverse is true for the uncertainty in *K* (Figure 4(c) and (f)). This behaviour is observed regardless of whether the Fisher–KPP model (Figure 4(a)–(c)) or the Porous Fisher model (Figure 4(d)–(f)) is assumed. Further insight is obtained through the posterior correlation coefficient matrix (see Appendix D, Table D.4), as *λ* and *K* are negatively correlated for all initial densities. This result is consistent with the results of Warne et al. (2017) and demonstrates that different experimental designs may be required to target different parameters. Lower initial densities result in data that only captures transient dynamics which maximises information related to *λ*. On the other hand, higher initial densities enable more precise estimation of limiting dynamics, that is, the effect of *K* can be observed. This feature would be difficult to elicit using traditional MLE-based methods.

The modes of the posterior marginal PDFs for *λ* and *K* are in agreement across both models (Figure 4(b)–(c) and (e)–(f)). However, the Porous Fisher model leads to lower uncertainty than the Fisher–KPP model in both of these parameters, regardless of initial density. This indicates that the uncertainty in the diffusion parameter, *D*_0_, has more of an impact on the other parameters under the Fisher–KPP model.

Comparing the posterior marginal PDFs for *D*_0_ requires some care. An initial inspection reveals the uncertainty in *D*_0_ looks significantly larger in the Porous Fisher model (Figure 4a) compared with the Fisher–KPP model (Figure 4d). However, the role of *D*_0_ in the density dependent diffusion of the Porous Fisher model (Equation (5)) is not the same as that in the Fickian diffusion of the Fisher–KPP model (Equation (4)). In the Fisher–KPP model, *D*_0_ is a constant cell diffusivity whereas in the Porous Fisher model *D*_0_ is the maximum diffusivity. Perhaps it would be more appropriate to give these two quantities different variables to make this point of distinction clear. Here, however, we have chosen to use the same variable to denote both quantities to be consistent with previous literature Jin et al. (2016b).

The most important aspect for the purposes of model comparison is the qualitative change in the diffusion parameter, *D*_0_, for different initial densities. For the Fisher–KPP model, not only does the uncertainty in *D*_0_ increase as the initial density increases, the mode also increases: *D*_0_ = 305.3 [*μ*m^2^/h] for low initial density; *D*_0_ = 850.9 [*μ*m^2^/h] for medium initial density; and *D*_0_ = 1371.4 [*μ*m^2^/h] for high initial density (Figure 4a). This is a clear indication that *D*_0_ depends on cell density, and this observation directly contradicts the implicit assumption made in invoking the Fisher–KPP model which treats *D*_0_ as a constant. This analysis would indicate that PC-3 cells exhibit density dependent motility where the diffusivity increases with the density. In contrast, the posterior marginal PDFs of *D*_0_ under the Porous Fisher model are very similar across initial densities (Figure 4d). Furthermore, the variance is consistent across all three initial densities considered. These results are in agreement with the observations of Jin et al. (2016b) who use the MLE to show that the Fisher–KPP model is inconsistent with the data. However, our Bayesian approach provides more detail through the reconstruction of the posterior PDF. Despite the fact that MLE model comparison selects the Fisher–KPP model as the preferred model, direct visualisation of the parameter uncertainty indicates otherwise.

### 4.3 Inference on the Generalised Porous Fisher model

The model comparison analysis performed in Section 4.2 is naturally extended by encoding model comparison as a single Bayesian inference problem applied to the Generalised Porous Fisher model (Equation (6)) with the exponent in the non-linear diffusion term, *r*, treated as an unknown parameter. This is essentially model comparison of a continuous population of models. This is another advantage of the Bayesian approach: model uncertainty can be treated identically to parameter uncertainty (Sunnåker et al., 2013).

The inference problem must now be slightly modified based on Section 4.2. The prior PDF becomes

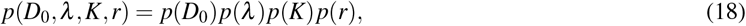

where

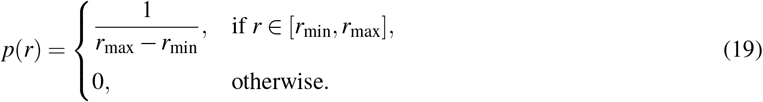

That is, *r ∼ 𝒰* (*r*_min_*, r*_max_). Equation (13), Equation (14) and Equation (15), respectively, specify *p*(*D*_0_), *p*(*λ*) and *p*(*K*). The limits that should be placed on *r* are unclear since *r* is has no physical interpretation and various values of *r* are used with little justification (Murray, 2002; Sherratt and Murray, 1990; Simpson et al., 2011). Since the most common values used in applications are *r* = 0 and *r* = 1, with the maximum known value used being *r* = 4, we take *r*_max_ = 8 so that we conservatively consider twice the largest value used in the mathematical biology literature. We also set *r*_min_ = *−*1 to allow for the possibility of *r <* 0, resulting in, so-called, “fast nonlinear diffusion” which is also thought to have some relevance to biological and ecological applications (King and McCabe, 2003).

Using the same assumptions for observation error as given in Section 4.2, the resulting likelihood is

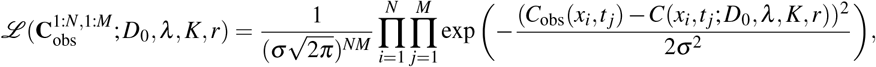

and the posterior PDF is

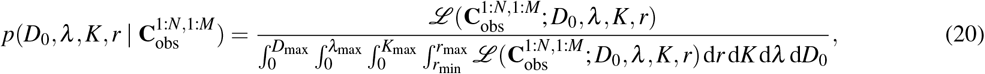

when [*D*_0_*, λ, K, r*] ∈ [0*, D*_max_]*×*[0*, λ*_max_]*×*[0*, K*_max_]*×*[*r*_min_*, r*_max_], otherwise 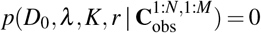. Here, *C*(*x_i_, t _j_*; *D*_0_*, λ, K, r*) is the numerical solution to the Generalised Porous Fisher equation, computed as per Appendix B. The posterior marginal PDFs are

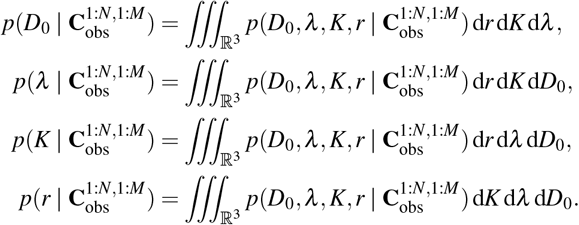

Computationally, low acceptance rates in the ABC rejection sampler render the technique ineffective for sampling this posterior distribution (Equation (20)). As a result, we apply an ABC variant of a Markov chain Monte Carlo (MCMC) sampler (Marjoram et al., 2003) (see Appendix C). While we find that the ABC MCMC sampler works well for this problem, other advanced ABC-based Monte Carlo schemes are also possible, such as sequential Monte Carlo (Sisson et al., 2007) and multilevel Monte Carlo (Warne et al., 2018). Using the ABC MCMC sampler, the four parameter posterior marginal PDFs derived from Equation (20) are estimated using the same data and observation error as in Section 4.2. Results are presented in Figure 5.

**Fig. 5.**
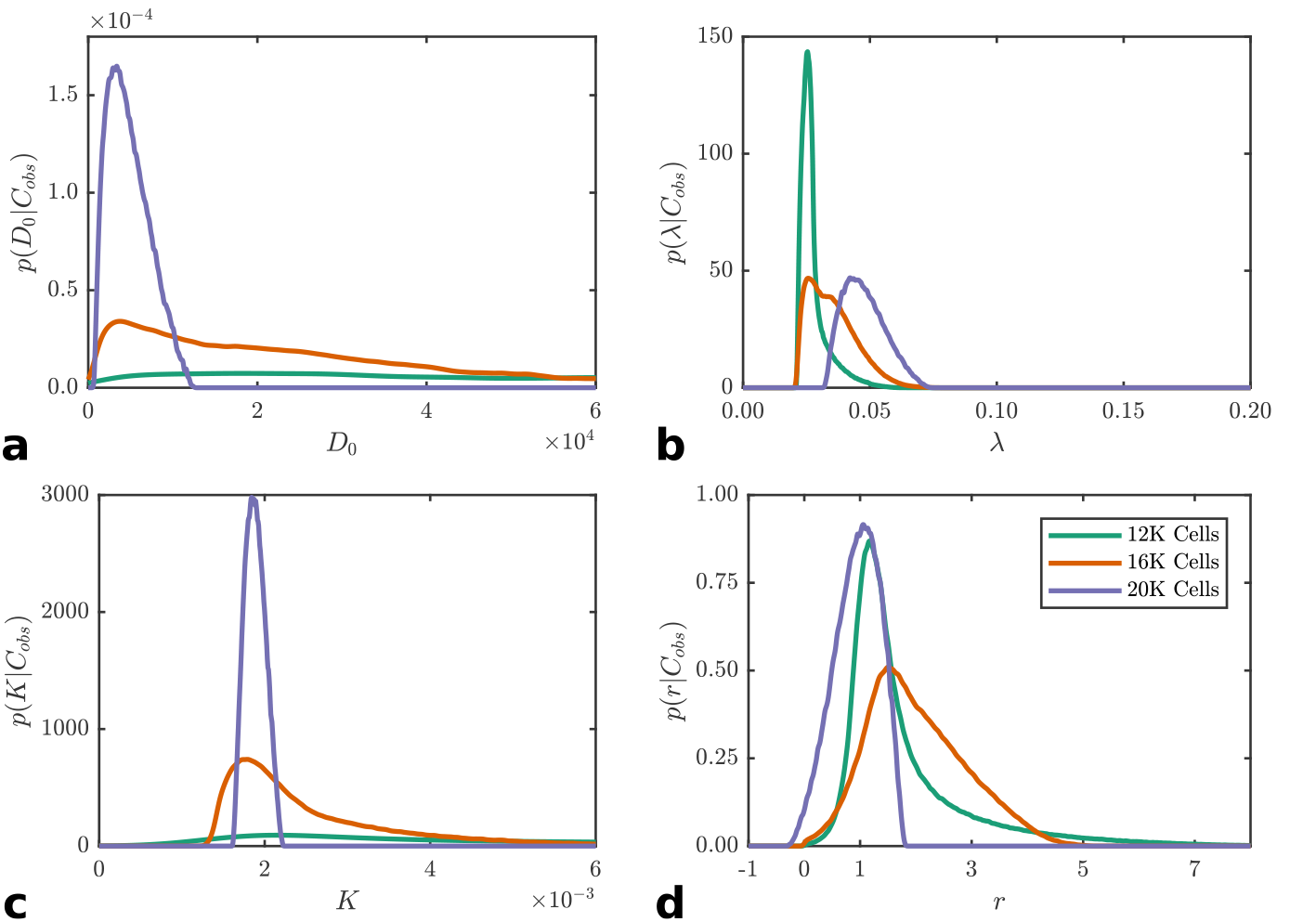
Marginal posterior probability densities obtained through Bayesian inference on the PC-3 scratch assay data under the Generalised Porous Fisher model. The spread in marginal densities demonstrate the degree of uncertainty in the diffusivity, *D*_0_, the proliferation rate, *λ*, the carrying capacity, *K*, and the exponent of the non-linear diffusion, *r*, under the three different initial conditions; 12, 000 cells (solid green), 16, 000 cells (solid orange), 20, 000 cells (solid purple).

The marginal posterior PDF for 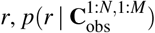, displays (Figure 5d) an overall higher probability density around *r* = 1 (corresponding to the Porous Fisher model) and very low probability density around *r* = 0 (corresponding to the Fisher–KPP model). However, this analysis also shows that other values of *r* may be justifiable. The uncertainty in *r* initially increases with increased initial density, but then decreases again with further increases in initial density. The same pattern occurs for the mode, as it transitions from *r ≈* 1 to *r ≈* 2 then back to *r ≈* 1.

The previously identified trade-off still exists between uncertainty in *K* and *λ* (Figure 5(b) and (c)) as the initial density increases. However, there is a qualitative difference in the marginal posterior PDFs compared to those in Figure 4(b),(c),(e) and (f). There is less consistency in the estimates across initial conditions, especially with the mode of *λ* apparently increasing as the initial density increases. For *D*_0_, almost no information is provided through the posterior marginal PDF except for data with high initial cell densities. Further analysis of the multivariate posterior marginal PDFs or the full joint posterior PDF would be required to obtain more information for this purpose. Such analysis is significantly more complex, and it may still yield a large degree of uncertainty in *D*_0_. In Appendix D an analysis is performed using bivariate marginal PDFs. It is clear the interactions between *r* and the other parameters are quite complex (see Figure D.2). Furthermore, as seen in Table D.4, *r* and *λ* are positively correlated for low initial density, but negatively correlated for high initial density; similarly, *r* and *K* are negatively correlated for low initial density, but positively correlated for high initial density. This kind of behaviour is difficult to interpret biologically, and we conclude that it is an artifact of using an overly complex model.

We demonstrate that a full Bayesian approach can easily incorporate comparison of a population of models. However, significantly more detailed analysis is required to interpret the results. Conversely, comparison of two distinct models though individual Bayesian inferences resulted in reasonable conclusions with minimal detailed analysis. A generalised model will never provide an lower fit in MLE (Stoica and Selen, 2004), however, there must be a point where the improved MLE is negligible (or non-existent) compared with the overall increase in uncertainty that must come with the generalisation. This raises the question whether the increased information obtained through a generalised model is worth the additional complexity.

## 5 Information criteria to balance model fit and complexity

In Sections 4.2 and 4.3, we observe that information gains from model generalisations may come at the cost of increased complexity and parameter uncertainty. We demonstrate in Section 3 and 4.2 that the definition of a good model must be based on more than the MLE alone. The theoretical underpinnings, explanatory power, biological feasibility, verifiability and complexity of a model are all important factors to consider when performing model selection (Box, 1976; Jin et al., 2016b; Sarapata and de Pillis, 2014; Slezak et al., 2010; Spiegelhalter et al., 2002). Overparameterised, complex models may fit the data well, however, the principle of Occam’s razor dictates that a simple model should be preferred wherever possible. In this section, we demonstrate the use of statistical measures, known as information criteria, that are designed to deal with trade-off between complexity and model fit (Gelman et al., 2014; Johnson and Omland, 2004).

### 5.1 Information criteria

Information criteria can be considered as methods for the ranking of models. Of the wide variety of information criteria available, many are based on rewarding models for lower residual error and penalising models for parameterisation (Gelman et al., 2014). Most information criteria are derived from decision theory and are based on the minimisation of some measure of information loss (Gelman et al., 2014; Johnson and Omland, 2004; Stoica and Selen, 2004). The resulting criteria, under suitable assumptions, are asymptotically proportional to information loss relative to an unknown true model (Gelman et al., 2014). That is, the model with the lowest information criterion value has the lowest information loss asymptotically. This is often taken to be a superior model from a decision theoretic perspective (Yang, 2005). However, caution must be taken when applying such measures because they are only guaranteed to inform correct decisions in the large sample limit (Gelman et al., 2014).

The three most fundamental information criteria are considered here, each of which have district properties, advantages and disadvantages. These information criteria have seen use in a limited set of biological applications (Johnson and Omland, 2004), and we are unaware of any application of these measures in the study of collective cell migration. It is important to note that there are many variants of the aforementioned criteria, however, we restrict ourselves the standard formulations in this work; for further information, see Gelman et al. (2014). We compare and contrast the information criteria results for the scratch assay data against the full Bayesian analysis performed in Section 4.

### 5.2 The Akaike information criterion

Akaike (1974) was the first to propose a solution to the key problem with MLE-based approaches, that is, overparameterised models will always be preferred (Akaike, 1974). Akaike considered the Kullback–Leibler (KL) information measure (Gelman et al., 2014; Kullback and Leibler, 1951),

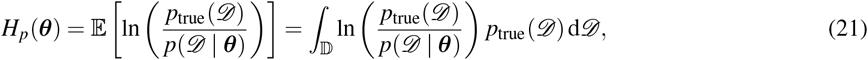

where ln(*⋅*) denotes the natural logarithm. The KL information measures the information loss, in the Shannon entropy sense (Shannon, 1948), incurred by assuming a model, *p*(*𝒟 |* ***θ***), of the data, *𝒟* ∈ 𝔻, instead of the hypothetical true model *p*_true_(*𝒟*) ^1^. Therefore, given two candidate models, *p*(*𝒟 |* ***θ***) and *q*(*𝒟 |* ***θ***), comparing *H_p_*(***θ***) and *H_q_*(***θ***) would reveal the model that minimises information loss. However, *p*_true_(*𝒟*) is unknown, so it is not possible to evaluate Equation (21) in practice. By analysing an asymptotic expansion of the KL information about the true parameter set, Akaike (1974) derives a penalty on the MLE based on the number of parameters that must be estimated. The result is the Akaike information criterion (AIC),

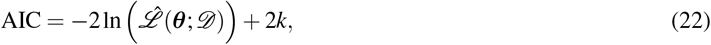

where *k* is the dimensionality of ***θ*** and *ℒ ̂*(***θ***; *𝒟*) is the maximum likelihood estimate, that is,

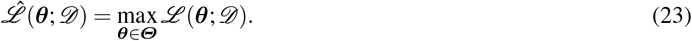

Due to the penalty incurred by the number of model parameters, a more complex model must improve the agreement with the data sufficiently to outperform a simpler model with an inferior maximum likelihood estimate. However, for models with the same number of parameters, the AIC is equivalent to the maximum likelihood estimate. When models have different numbers of parameters, the AIC favours simpler models. Unfortunately, the AIC is not an asymptotically consistent estimator, that is, as *n →* ∞, the AIC is not guaranteed to converge to a unique model (Yang, 2005). However, the AIC will select the model with optimal residual error, since it may be viewed as the MLE with a bias correction to compensate for overfitting (Yang, 2005).

### 5.3 The Bayesian information criterion

An alternative approach, the Bayesian information criterion (BIC), has different theoretical foundations (Schwarz, 1978). Schwarz (1978) considered Bayes estimators instead of the KL information, thereby defining the model with the maximum *a posteriori* probability to be the optimal choice. The resulting BIC is,

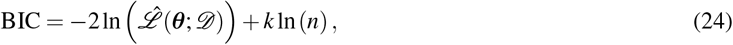

where *k* is the dimensionality of ***θ***, *n* is the dimensionality of *𝒟* and *ℒ ̂*(***θ***; *𝒟*) is the maximum likelihood estimate as given in Equation (23). The BIC is a consistent approximation to the maximum *a posteriori* estimate (i.e., the mode of the posterior PDF) and is independent of model priors provided *k/n* ≪ 1. Compared with the AIC, the BIC attributes a larger penalty for model complexity when *n ≥* 8, thus the BIC favours simplicity more than the AIC. The BIC also has the advantage of being consistent as *n →* ∞. In addition, the BIC is asymtotically equivalent to the comparison of Bayes factors, which are considered more correct by some (Pooley and Marion, 2018), however, others note that the usage of Bayes factors assumes that the model prior covers the correct model (Gelman et al., 2004). The BIC, however, will not always select the model with the optimal residual error compared with the AIC (Yang, 2005).

### 5.4 The deviance information criterion

The prevalence of Monte Carlo sampling in practical Bayesian applications was the motivation for Spiegelhalter et al. (2002) to develop the deviance information criterion (DIC). The DIC has particular computational advantages for MCMC sampling (Gelman et al., 2014; Spiegelhalter et al., 2002). Like the AIC, the DIC has its theoretical foundation in minimsation of the KL information loss (Gelman et al., 2004). However, unlike the AIC and BIC, the DIC does not require the maximum likelihood estimate to be computed, but is based on expectation calculations. As a result, the DIC is ideal for Monte Carlo integration schemes (Gelman et al., 2014). The DIC is given by,

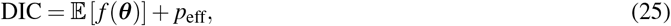

where *f* (*⋅*) is the deviance function,

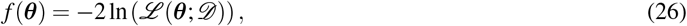

and *p*_eff_ is the effective number of parameters,

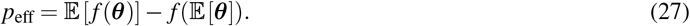

Importantly, the expectations are evaluated with respect to the posterior probability measure. The DIC is conceptually quite different to the AIC and BIC. Firstly, the DIC uses an averaged likelihood rather than point estimates like the AIC and BIC. Secondly, the DIC effective parameter term, *p*_eff_, attempts to distinguish between information obtained through the prior distribution rather than the data (Gelman et al., 2004).

Because the averaged likelihood is utilised, it can be considered more closely aligned with a Bayesian viewpoint that aims to use information from the entire posterior distribution (Gelman et al., 2014). However, the use of the DIC as a reliable information criterion has been debated in the literature, in particular there is concern that the DIC is inconsistent with Bayes factors, that is, the DIC may fail to select the true model even if it is among the set of candidates (Pooley and Marion, 2018; Spiegelhalter et al., 2014). However, due to simplicity of calculation via Monte Carlo integration and applicability to hierarchical models, the DIC has been widely adopted for practical applications (Gelman et al., 2014; Spiegelhalter et al., 2014).

### 5.5 Evaluation of continuum models using information criteria

We compute the AIC, BIC and DIC for the Fisher–KPP, Porous Fisher and Generalised Porous Fisher models given the data derived from Jin et al. (2016b). The results are presented for each initial condition in Table 1.

**Table 1.** Information criteria for initial conditions of 12, 000 cells, 16, 000 cells and 20, 000 cells.

| Model | 12,000 cells |  |  | 16,000 cells |  |  | 20,000 cells |  |  |
| --- | --- | --- | --- | --- | --- | --- | --- | --- | --- |
|  | DIC | BIC | AIC | DIC | BIC | AIC | DIC | BIC | AIC |
| Fisher–KPP | -2386.08 | -2377.06 | -2386.13 | -2367.03 | -2358.00 | -2367.07 | -2268.60 | -2259.54 | -2268.61 |
| Porous Fisher | -2398.30 | -2389.87 | -2398.94 | -2392.87 | -2383.59 | -2392.66 | -2278.58 | -2269.41 | -2278.49 |
| Generalised Porous Fisher | -2399.07 | -2387.92 | -2400.01 | -2390.88 | -2378.84 | -2390.93 | -2279.36 | -2267.59 | -2279.69 |

Across all initial conditions, the BIC consistently selects the Porous Fisher model. The consistency of the BIC is expected due to its theoretical basis of Bayes estimators (Schwarz, 1978; Spiegelhalter et al., 2014; Yang, 2005), that is, if the true model was in the set of candidates, then BIC would asymptotically select that model. The BIC results are also consistent with the full Bayesian analysis performed in Section 4.

The AIC and DIC are in very close agreement, preferring the Generalised Porous Fisher model for both low and high density initial conditions (Table 1), but favouring the Porous Fisher Model for the intermediate initial density. The agreement between the AIC and DIC is also expected theoretically since they are both derived from the KL information loss. In fact, if 𝔼 [*f* (***θ***)] *≈ −*2 ln *ℒ ̂* (***θ***; *𝒟*) and *p*_eff_ *≈ k* then Equation (22) and Equation (25) are approximately equivalent. The results in Table 1 indicate these relationships likely hold in our case.

Overall, the information criteria provide clear indications that the Fisher–KPP model does not sufficiently describe the collective behaviour of the PC-3 cells for any initial density. In Table 1 we see there is good agreement between the rankings suggested by the AIC and DIC, and some disagreement in the ranking suggested by the BIC. However, the improvement in the AIC and DIC for the Generalised Porous Fisher model over the Porous Fisher model is negligible compared with the improvement in the AIC and DIC for the Porous Fisher model over Fisher–KPP model. Furthermore, the BIC consistently selects the Porous Fisher model for each initial density. Therefore, we conclude that the Porous Fisher model represents the best trade-off between model fit and complexity.

## 6 Discussion and outlook

We have demonstrated in Section 3 that traditional MLE methods of model calibration and model selection do not provide a satisfactory method to compare continuum models of collective cell spreading and proliferation since MLE methods favour overparameterised models. The Bayesian approach presented in Section 4 and the analysis using information criteria that is presented in Section 5 provide a significantly more robust methodology to evaluate the ability of continuum models to explain collective cell behaviour. This methodology has enabled us to present a clear example of when model generalisation leads to less consistent and more uncertain parameter estimation. The Bayesian analysis presented in Section 4 indicates that the Porous Fisher model provides the most consistent parameter estimates, and information criteria demonstrated in Section 5 suggest that the Porous Fisher model represents the optimal trade-off between model fit and complexity.

Information criteria provide an objective approach to model selection that take into account model fitness and complexity. However, reducing model comparison down to a single scalar comparison is bound to disregard some important aspects of the model comparison problem (Gelman et al., 2014; Spiegelhalter et al., 2014). On the other hand, Bayesian posterior distributions provide a rich source of information. For example, the shifts in modes and support in the diffusion parameter, *D*_0_, for both the Fisher–KPP model (Figure 4a) and Generalised Porous Fisher model (Figure 5a) are not directly identifiable using information criteria. If an objective decision rule is required, then a clear understanding on the assumptions and asymptotic properties of the criterion used is essential. We conclude, along with Yang (2005), that the BIC is most appropriate to select the best overall model, whereas the AIC and DIC are better suited if the goal is prediction over a short time-scale. Bayes factors are another alternative, however, their results can be highly sensitive to the choice of model prior distributions, especially when model parameters are continuous (Gelman et al., 2014; Pooley and Marion, 2018).

The Bayesian approach we take in Section 4 is the most direct and intuitive approach to model selection. Model inconsistency across data sets is apparent, through comparing marginal posterior PDFs for different datasets. Parameters that the data provides little information about are also highlighted this way. Of course, there are many extensions to the analysis that could be performed. The correlation structures of the full joint posterior PDF can elicit connections between model parameters that cannot be identified from the marginal posterior PDFs alone. Visualisation of bivariate marginal posterior PDFs are also useful to see more details on these interactions. We have provided extended results in Appendix D that were excluded from the main manuscript for clarity in our key points: 1) increasing model complexity increases uncertainty in parameter estimates; and 2) model consistency must be evaluated using multiple datasets with different initial conditions.

Jin et al. (2016b) and Warne et al. (2017) highlight the importance of modelling the uncertainty in the initial density of cell culture assays. In particular, Jin et al. (2016b) demonstrate that variability in the initial density is not negligible across identically prepared replicates. Warne et al. (2017) show that this variation, if properly modelled as a random variable, greatly impacts the uncertainty in the estimates of carrying capacity, *K*. We extend this analysis in the context of the continuum models considered in this work and include results in Appendix D. The key result is that parameter uncertainty is amplified in this more realistic, but rarely considered, case.

While we have primarily focused on model selection across different cell motility mechanisms, others have proposed models including generalisations of the source term in Equation (1) (Browning et al., 2017; Jin et al., 2016a; Tsoularis and Wallace, 2002), the inclusion of growth factors (Jin et al., 2016b; Sherratt and Murray, 1990), or chemotaxis (Bianchi et al., 2016). Such extensions are of interest and should be the subject of future research. We do not specifically investigate these here, though our analysis can be repeated in such cases at an increased computational expense. However, given the significant increase in parameter uncertainty incurred by the Generalised Porous Fisher model, that included only a fourth parameter, it is highly likely that scratch assay data is insufficient to provide any model certainty for these more complex extensions involving many more parameters. For example, as discussed by Johnston et al. (2015), chemotaxis amounts to setting **J**(**x***, t*) = *−D*(*C*(**x***, t*))∇*C*(**x***, t*) + *χC*(**x***, t*)∇*G*(**x***, t*) in Equation (1), leading to

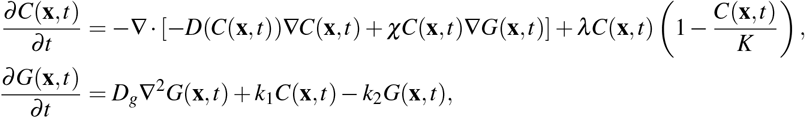

where *χ* is the chemotatic sensitivity coefficient, *G*(**x***, t*) *>* 0 is the concentration of a diffusive chemical signal, *D_g_ >* 0 is the diffusivity of the chemical, *k*_1_ *>* 0 and *k*_2_ *>* 0 are kinetic rate parameters for chemical production (by cells) and degradation, respectively. If *χ <* 0, then cells are repelled by the diffusive signal, and if *χ >* 0, then cells are attracted to it. Therefore, we have a minimum of seven parameters, with ***θ*** = [*D*_0_*, λ, K, χ, D_g_, k*_1_*, k*_2_]. Furthermore, the model includes a single chemical species only and a realistic model would need to account more many interacting chemical factors. Standard experimental protocols of cell culture assays do not measure this information. Cell density data alone will not be sufficient to calibrate this model without significant levels of uncertainty. Increased number of parameters impacts the convergence time of MCMC samples (Gelman et al., 2014).

We do not advocate against the validity or utility of more complex models of collective cell motility and proliferation. There are many biologically based rationales for including extra biophysical and biochemical factors in a given model (Bianchi et al., 2016; Nardini et al., 2016). However, our results do indicate that current *in vitro* cell culture assay data are not informative enough to distinguish between these models in practice, a point that is rarely discussed in the literature. This work is intended to motivate more detailed, Bayesian model selection within the mathematical biology community, and provide evidence that higher quality experimental methods and image analysis tools are required to validate and compare the biological hypotheses of the future.

Our results have broad implications for the mathematical biology community. Specifically, if a complex model is to be applied, then sufficient data must be collected in order to produce meaningful calibrations. Studies that compare hypotheses should also take model complexity and parameter uncertainty into account when making conclusions. The Bayesian framework, as presented here, provides tools that are designed to assist in these aspects. We suggest such techniques should be widely adopted.

## Acknowledgements

This work is supported by the Australian Research Council (DP170100474). Ruth E. Baker is a Royal Society Wolfson Research Merit Award holder and a Leverhulme Research Fellow, and also acknowledges the Biotechnology and Biological Sciences Research Council for funding via grant no. BB/R000816/1. Computational resources were provide by the eResearch Office, Queensland University of Technology.

## Appendix A Analysis of wave front concavity

The Generalised Porous Fisher model (Equation (6) in the main text) with *λ* = 0, in one dimension is

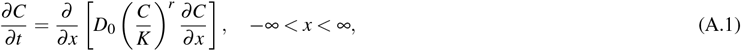

where *D*_0_ is the free diffusivity and *K* the cell carrying capacity density. For the initial condition *C*(*x*, 0) = *C*_0_*δ* (*x*), Equation (A.1) has an exact solution,

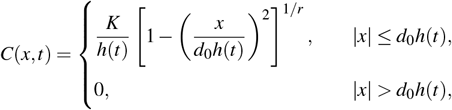

where 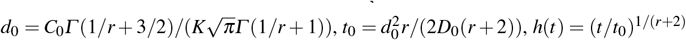 and *Γ* (*x*) is the Gamma function. This solution, often called the source solution for the porous media equation, has compact support, *x* ∈ [*−d*_0_*h*(*t*)*, d*_0_*h*(*t*)]. Here, *|x|* = *d*_0_*h*(*t*) are the contact points. This solution is very different to the source solution for the linear diffusion equation, *r* = 0, which is a Gaussian function without compact support (Barenblatt, 2003; Crank, 1975).

Without loss of generality, we now only consider the positive real line *x ≥* 0. The cell density is always decreasing as we approach the contact point, that is, *∂C/∂ x <* 0 for 0 *< x < d*_0_*h*(*t*). Specifically, we have

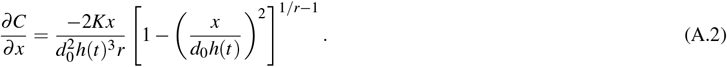

From Equation (A.2) three different front properties are possible. As *x → d*_0_*h*(*t*) we observe: (i) a smooth front, for 0 *< r <* 1, as in Figure 2e– (h), with *∂C/∂ x →* 0; (ii) a sharp front with finite negative slope, for *r* = 1, as in Figure 2(i)–(l), with 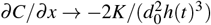; and (iii) a sharp front with unbounded negative slope, for *r >* 1, as in Figure 2(m)–(p), with *∂C/∂ x → −*∞.

To explore the concavity of the density profile, *C*(*x, t*), at the contact point, it is sufficient to show explore how the sign of *∂* ^2^*C/∂ x*^2^ at the contact point depends on *r*. The second derivative with respect to *x*, for *r >* 0, is

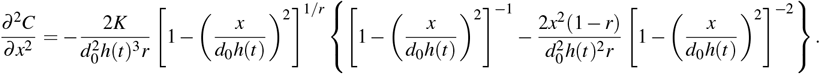

We have, for 0 *< r <* 1, that *∂* ^2^*C/∂ x*^2^ *>* 0 as *x → d*_0_*h*(*t*). For *r ≥* 1, *∂*^2^*C/∂ x*^2^ *<* 0 as *x → d*_0_*h*(*t*). Hence, at the contact point, *x* = *d*_0_*h*(*t*), the solution is concave down for *r ≥* 1, and concave up otherwise.

## Appendix B Numerical scheme

Here we describe our numerical scheme for the computational solution to the following reaction–diffusion equation:

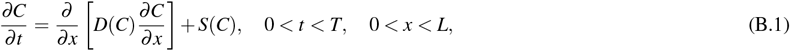

with initial condition,

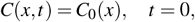

and boundary conditions,

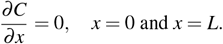

Consider *N* points in space, 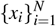, with *x*_1_ = 0, *x_N_* = *L* and *∆ x* = *x_i_*_+1_ *− x_i_* for all *i* = [1, 2*,…, N*]. Similarly, define *M* temporal points, 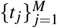, with *t*_1_ = 0, *t_M_* = *T* and *∆t* = *t _j_*_+1_ *− t _j_* for all *j* = [1, 2*,…, T*]. Next, define the notation, *C_i_*_+*k*_ = *C*(*x_i_* + *k∆ x, t*), and 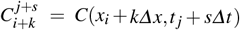.

Let *J*(*C*) = *−D*(*C*)*∂C/∂ x* and substitute into Equation (B.1) to yield

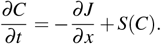

At the *i*th point, apply a first order central difference to *∂ J/∂ x* with step *∆ x/*2. The result is the system of ODEs

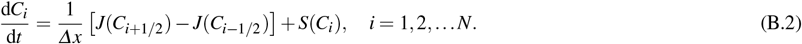

Similarly, a first order central difference is applied to *J*(*C_i_*_+1*/*2_) and *J*(*C_i−_*_1*/*2_) using the step *∆ x/*2 yields

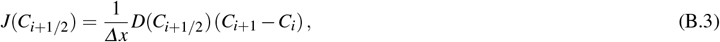

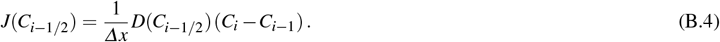

It is important to note that we will only obtain a solution for *C_i_*_+*k*_ at integer values of *k*, therefore the evaluation of the diffusion terms in Equation (B.3) and Equation (B.4) cannot be directly computed since *k* = *±*1*/*2. We thus approximate with a linear interpolation,

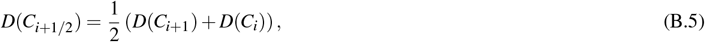

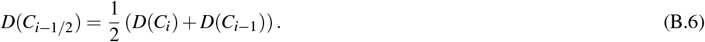

After substitution of Equation (B.3), Equation (B.4), Equation (B.5) and Equation (B.6) into Equation (B.2), we have the coupled system of non-linear ODEs defined in terms of our spatial discretisation,

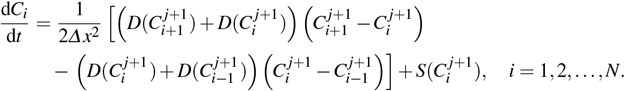

The no-flux boundaries are enforced using first order forward differences

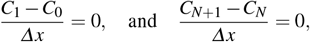

where *C*_0_ and *C_N_*_+1_ represent the solution at “ghost nodes” that are not a part of the domain.

The ODEs are discretised in time using a first order backward difference method leading to the backward-time, centered-space (BTCS) scheme,

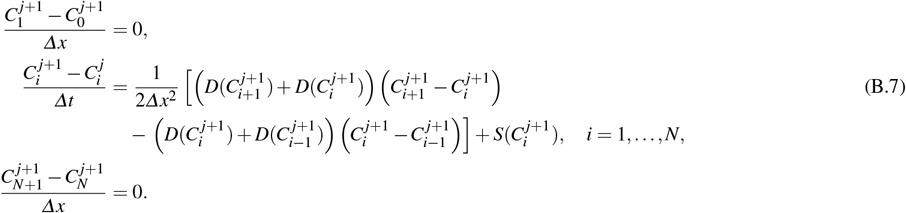

While this scheme is first order in time and space, it has the advantage of unconditional stability.

Since the scheme is implicit, a non-linear root finding solver is required to compute solution at *t _j_*_+1_ given a previously computed solution at time *t _j_*. To achieve this we apply fixed-point iteration. We re-arrange the system to be of the form **C** ^*j*+1^ = **G**(**C^j^**^+**1**^) where 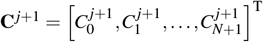. That is,

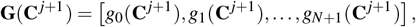

where

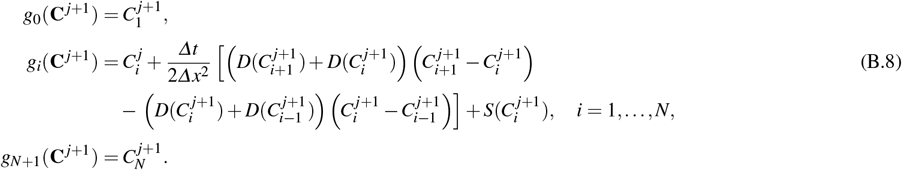

We then define the sequence {**X**^*k*^}_*k≥*0_, generated through the non-linear recurrence relation **X**^*k*+1^ = **G**(**X**^*k*^) with **X**^0^ = **C**^*j*^. This sequence is iterated until *||***X**^*k*+1^ *−* **X**^*k*^|| *< τ*, where *τ* is the error tolerance and *||⋅||*_2_ is the Euclidean vector norm. Once the sequence has converged, we set **C** ^*j*+1^ = **X**^*k*+1^ and continue to solve for the next time step.

For a given set of model parameters, the spatial and temporal step sizes, *∆ x* and *∆t*, need to be selected. In particular, the following condition must hold to ensure accuracy, max_*C*∈[0*,K*]_ *D*(*C*) *< ∆ x*^2^*/∆t*. We then refine *∆ x* and *∆t* together to ensure solutions are independent of the discretisation. Note that as *r* increases, higher values of *D*(*C*) become valid, therefore particular attention is required to generate Figure 5 in the main text. The values of *∆ x*, *∆t* and *τ* used for the simulations in this work are shown in Table B.1. Note, that is all cases the discretisation is more refined than required to solve the given problem accurately.

**Table B.1.** Discretisation and tolerance for numerical simulations.

| Application | $\Delta x$ [$\mu\text{m}$] | $\Delta t$ [h] | $\tau$ |
| --- | --- | --- | --- |
| Example evolutions (Figure 2) | 12.80 | $6.0 \times 10^{-3}$ | $1.0 \times 10^{-4}$ |
| Inference on Fisher–KPP (Figure 4) | 9.95 | $3.0 \times 10^{-3}$ | $1.0 \times 10^{-6}$ |
| Inference on Porous Fisher (Figure 4) | 9.95 | $1.5 \times 10^{-3}$ | $1.0 \times 10^{-6}$ |
| Inference on Generalised Porous Fisher (Figure 5) | 8.22 | $7.5 \times 10^{-4}$ | $1.0 \times 10^{-6}$ |

## Appendix C Computational inference

The Bayesian inference problems described in the main text all require the computation of the posterior PDF. Up to a normalisation constant, the posterior PDF is given by

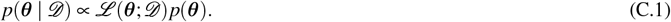

If the posterior distribution can be sampled, the posterior PDF may be determined by using Monte Carlo integration. Thus, the main requirement is a method of generating *N* independent, identically distributed (i.i.d.) samples from the posterior distribution.

For many applications of practical interest, Equation (C.1) cannot be used directly to generate the samples required since the likelihood is often intractable. Approximate Bayesian computation (ABC) techniques resolve this complexity through the approximation (Sunnåker et al., 2013)

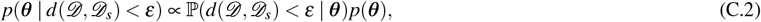

where *d*(*𝒟, 𝒟_s_*) is a discrepancy metric between the true data, *𝒟*, and simulated data, *𝒟_s_ ∼ ℒ* (***θ***; *𝒟_s_*) and *ε* is the discrepancy threshold. ABC methods have the property that *p*(***θ*** *| d*(*𝒟, 𝒟_s_*) *< ε*) *→ p*(***θ*** *| 𝒟*) as *ε →* 0. This leads directly to the ABC rejection sampling algorithm (Algorithm C.1). For deterministic models, under the assumption of Gaussian observation errors, *ε/σ* ≪ 1, and *d*(*𝒟, 𝒟_s_*) taken as the sum of the squared errors, it can be shown that ABC methods are equivalent to exact posterior sampling (Wilkinson, 2013).

### Algorithm C.1 ABC rejection sampling

~~~
1: **for** *i* = 1*,…, N* **do**
2:    **repeat**
3:      Sample prior, ***θ***^*^ *∼ p*(***θ***).
4:      Generate data, *𝒟_s_ ∼ ℒ* (***θ***^*^; *𝒟_s_*).
5:    **until** *d*(*𝒟, 𝒟_s_*) *≤ ε*
6:    Set ***θ***^*i*^ ← ***θ***^*^.
7: **end for**
~~~

In some cases, the acceptance probability in Algorithm C.1 is computationally prohibitive for small *ε*. In such situations, an ABC extension to Markov Chain Monte Carlo sampling may be applied (Marjoram et al., 2003). The resulting ABC MCMC sampling method (Algorithm C.2), under reasonable conditions on the proposal kernel *K*(***θ****^i^ |****θ***^*i−*1^), simulates a Markov Chain with *p*(***θ*** *| d*(*𝒟, 𝒟_s_*) *< ε*) (Equation (C.2)) as its stationary distribution. It is essential to simulate the Markov Chain for a sufficiently long time such that the *N_T_* dependent samples are effectively equivalent to the required *N* i.i.d. samples.

### Algorithm C.2 ABC Markov chain Monte Carlo sampling

~~~
1: Given initial sample ***θ***^1^ *∼ p*(***θ*** *| d*(*𝒟, 𝒟_s_*) *< ε*).
2: **for** *i* = 2*,…, N_T_* **do**
3:     Sample transition kernel, ***θ***^*^ *∼ K*(***θ*** *|* ***θ***^*i−*1^).
4:     Generate data, *𝒟_s_ ∼ ℒ* (***θ***^*^; *𝒟_s_*).
5.     **If** *d*(*𝒟, 𝒟_s_*) *≤ ε* **then**
6.        Set *h ←* min *p*(***θ***^*^)*K*(***θ***^*i−*1^ *|* ***θ***^*^)*/p*(***θ***^*i−*1^)*K*(***θ***^*^ *|* ***θ***^*i−*1^), 1).
7:        Sample uniform distribution, *u ∼ 𝒰* (0, 1).
8.        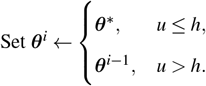
9:     **else**
10:       Set ***θ***^*i*^ ← ***θ***^*i−*1^.
11:    **end if**
12: **end for**
~~~

Using either ABC rejection sampling (Algorithm C.1) or ABC MCMC sampling (Algorithm C.2), we can apply Monte Carlo integration to compute the posterior PDF as given in Equation (C.2). For simplicity, we focus on the approximation of the *j*th marginal posterior PDF (Silverman, 1986),

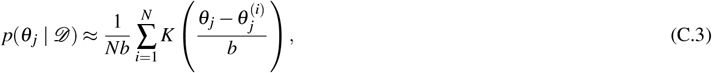

where *θ _j_* is the *j*th element of ***θ***, 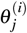 are the *j*th elements of 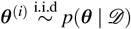, *b* is the smoothing parameter and *K*(*x*) is the smoothing kernel with property 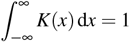.

## Appendix D Additional results

In this appendix, we present extended results that are excluded from the main text for brevity. We provide more detailed information on the Bayesian analysis presented in Sections 4.2 and 4.3. Furthermore, we extend the Bayesian inference problem, as provided in Section 4.2, to account for the treatment of uncertainty in the initial condition.

### D.1 Joint posterior features

Here we report various descriptive statistics for the joint posterior PDFs computed in Section 4. For each posterior distribution we report the posterior mode, the posterior mean, the variance/covariance matrix and the correlation coefficient matrix.

Given cell density data, *𝓓*, a set of continuum model parameters, ***θ***, in parameter space ***Θ*** ⊆ ℝ^*k*^ with *k >* 0, and a model implied through a likelihood function, *ℒ* (***θ***; *𝓓*), then summary statistics can be computed from the joint posterior, *p*(***θ*** *| 𝓓*), to obtain estimates and uncertainties on the true parameters. The maximum *a posteriori* (MAP) parameter estimate is the parameter set with the greatest posterior probability density as given by the posterior mode,

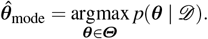

The posterior mean is the central tendency of the parameters,

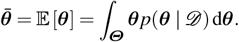

The variance/covariance, matrix *Σ ∈* R^*k×k*^, provides information on the multivariate uncertainties, that is the spread of parameters. The (*i, j*)th element of *Σ*, denoted by *σ_i, j_*, is given by

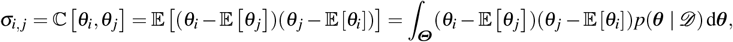

where *θ_i_* and *θ_j_* are the *i*th and *j*th elements of ***θ***. Note that ℂ [*θ_i_, θ_i_*] = 𝕍 [*θ_i_*] and *σ_i, j_* = *σ_j,i_*. Lastly, the correlation coefficient matrix *R* ∈ R^*k×k*^ measures the linear dependence between parameter pairs. The (*i, j*)th element of *R*, denoted by *ρ_i, j_*, is given by

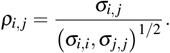

Note *ρ_i,i_* = 1 for all *i* ∈ [1*, k*], and *ρ_i, j_* = *ρ _j,i_*. The results of all these statistics, for the inference problems considered in the main text, are presented in tables D.1, D.2, D.3, and D.4.

**Table D.1.** MAP parameter estimates (posterior modes) from posterior PDFs using initial conditions of 12, 000 cells, 16, 000 cells and 20, 000 cells.

| Inference problem |  | MAP parameter estimate (posterior mode) |  |  |  |
| --- | --- | --- | --- | --- | --- |
| Model | Initial condition | $D_0$ [$\mu\text{m}^2/\text{h}$] | $\lambda$ [$1/\text{h}$] | $K$ [$\text{cells}/\mu\text{m}^2$] | $r$ [-] |
| Fisher-KPP | 12,000 | 325 | $3.05 \times 10^{-2}$ | $1.85 \times 10^{-3}$ | - |
| Fisher-KPP | 16,000 | 1,010 | $3.14 \times 10^{-2}$ | $1.49 \times 10^{-3}$ | - |
| Fisher-KPP | 20,000 | 1,354 | $3.56 \times 10^{-2}$ | $1.65 \times 10^{-3}$ | - |
| Porous Fisher | 12,000 | 3,425 | $3.17 \times 10^{-2}$ | $2.58 \times 10^{-3}$ | - |
| Porous Fisher | 16,000 | 3,134 | $3.77 \times 10^{-2}$ | $1.56 \times 10^{-3}$ | - |
| Porous Fisher | 20,000 | 4,933 | $5.32 \times 10^{-2}$ | $1.69 \times 10^{-3}$ | - |
| Generalised Porous Fisher | 12,000 | 7,857 | $2.66 \times 10^{-2}$ | $1.01 \times 10^{-2}$ | 1.37 |
| Generalised Porous Fisher | 16,000 | 90,234 | $2.43 \times 10^{-2}$ | $7.18 \times 10^{-3}$ | 1.58 |
| Generalised Porous Fisher | 20,000 | 11,990 | $3.46 \times 10^{-2}$ | $2.10 \times 10^{-3}$ | 1.65 |

**Table D.2.**
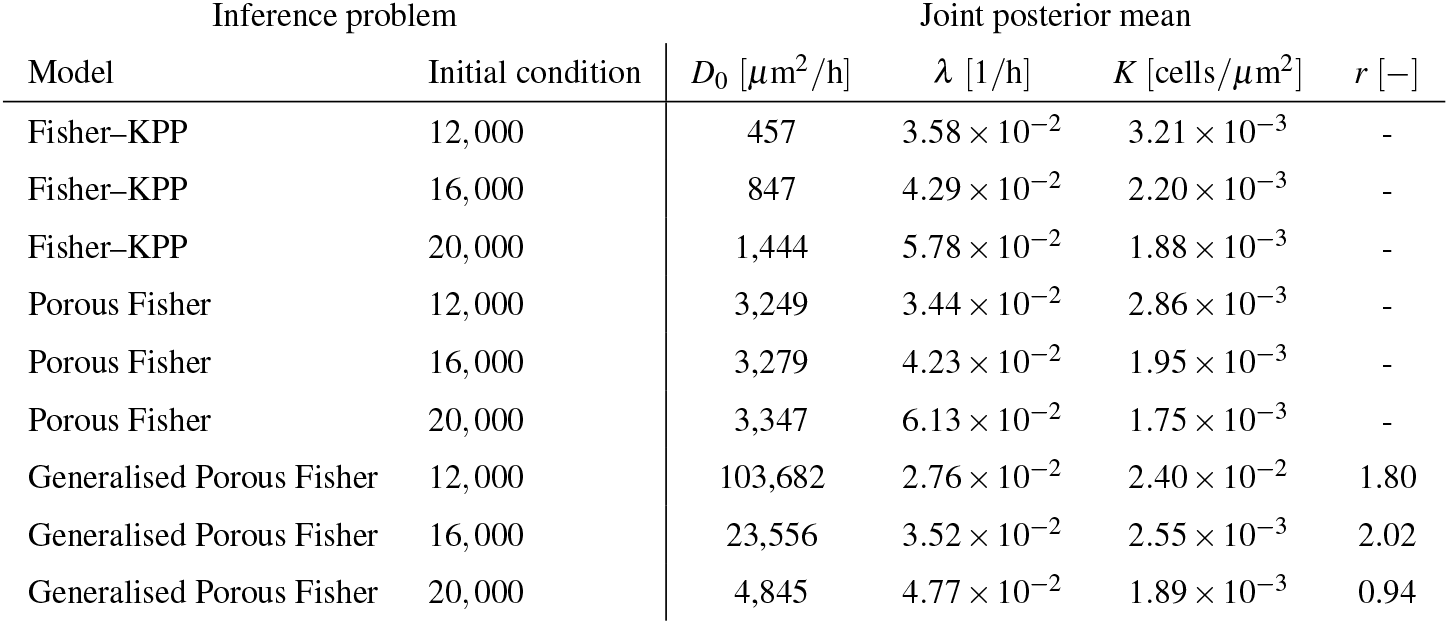
Joint posterior means for posterior PDFs using initial conditions of 12, 000 cells, 16, 000 cells and 20, 000 cells.

**Table D.3.**
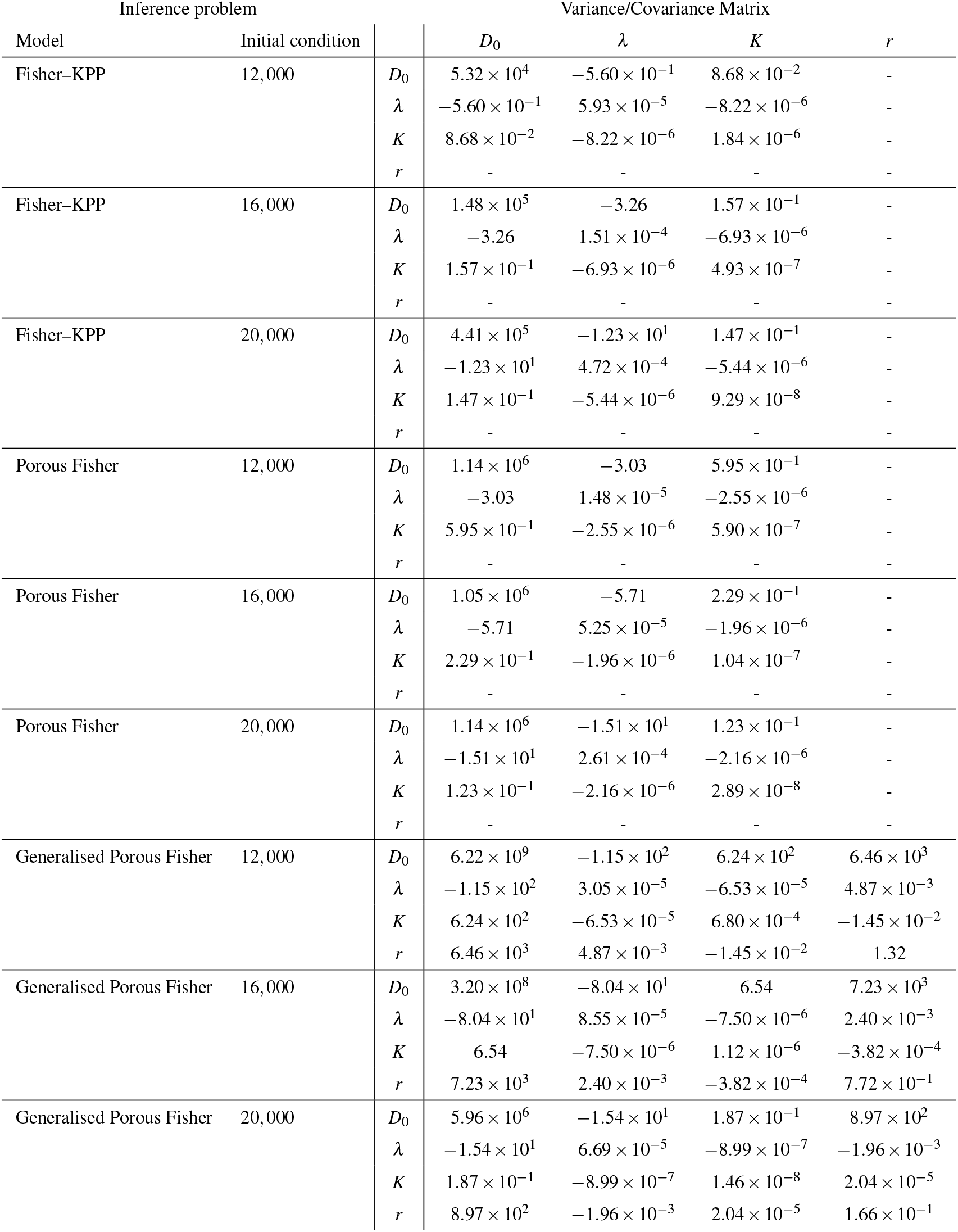
Variance/covariance matricies for posterior PDFs using initial conditions of 12, 000 cells, 16, 000 cells and 20, 000 cells.

**Table D.4.** Correlation coefficient matricies for posterior PDFs using initial conditions of 12, 000 cells, 16, 000 cells and 20, 000 cells.

| Inference problem |  | Correlation coefficient Matrix |  |  |  |  |
| --- | --- | --- | --- | --- | --- | --- |
| Model | Initial condition | | $D_0$ | $\lambda$ | $K$ | $r$ |
| Fisher-KPP | 12,000 | $D_0$ | 1.00 | $-3.15 \times 10^{-1}$ | $2.77 \times 10^{-1}$ | - |
| | | $\lambda$ | $-3.15 \times 10^{-1}$ | 1.00 | $-7.86 \times 10^{-1}$ | - |
| | | $K$ | $2.77 \times 10^{-1}$ | $-7.86 \times 10^{-1}$ | 1.00 | - |
| | | $r$ | - | - | - | - |
| Fisher-KPP | 16,000 | $D_0$ | 1.00 | $-6.90 \times 10^{-1}$ | $5.83 \times 10^{-1}$ | - |
| | | $\lambda$ | $-6.90 \times 10^{-1}$ | 1.00 | $-8.04 \times 10^{-1}$ | - |
| | | $K$ | $5.83 \times 10^{-1}$ | $-8.04 \times 10^{-1}$ | 1.00 | - |
| | | $r$ | - | - | - | - |
| Fisher-KPP | 20,000 | $D_0$ | 1.00 | $-8.50 \times 10^{-1}$ | $7.27 \times 10^{-1}$ | - |
| | | $\lambda$ | $-8.50 \times 10^{-1}$ | 1.00 | $-8.21 \times 10^{-1}$ | - |
| | | $K$ | $7.27 \times 10^{-1}$ | $-8.21 \times 10^{-1}$ | 1.00 | - |
| | | $r$ | - | - | - | - |
| Porous Fisher | 12,000 | $D_0$ | 1.00 | $-7.36 \times 10^{-1}$ | $7.25 \times 10^{-1}$ | - |
| | | $\lambda$ | $-7.36 \times 10^{-1}$ | 1.00 | $-8.61 \times 10^{-1}$ | - |
| | | $K$ | $7.25 \times 10^{-1}$ | $-8.61 \times 10^{-1}$ | 1.00 | - |
| | | $r$ | - | - | - | - |
| Porous Fisher | 16,000 | $D_0$ | 1.00 | $-7.68 \times 10^{-1}$ | $6.90 \times 10^{-1}$ | - |
| | | $\lambda$ | $-7.68 \times 10^{-1}$ | 1.00 | $-8.36 \times 10^{-1}$ | - |
| | | $K$ | $6.90 \times 10^{-1}$ | $-8.36 \times 10^{-1}$ | 1.00 | - |
| | | $r$ | - | - | - | - |
| Porous Fisher | 20,000 | $D_0$ | 1.00 | $-8.76 \times 10^{-1}$ | $6.80 \times 10^{-1}$ | - |
| | | $\lambda$ | $-8.76 \times 10^{-1}$ | 1.00 | $-7.87 \times 10^{-1}$ | - |
| | | $K$ | $6.80 \times 10^{-1}$ | $-7.87 \times 10^{-1}$ | 1.00 | - |
| | | $r$ | - | - | - | - |
| Generalised Porous Fisher | 12,000 | $D_0$ | 1.00 | $-2.64 \times 10^{-1}$ | $3.03 \times 10^{-1}$ | $7.12 \times 10^{-2}$ |
| | | $\lambda$ | $-2.64 \times 10^{-1}$ | 1.00 | $-4.53 \times 10^{-1}$ | $7.67 \times 10^{-1}$ |
| | | $K$ | $3.03 \times 10^{-1}$ | $-4.53 \times 10^{-1}$ | 1.00 | $-4.83 \times 10^{-1}$ |
| | | $r$ | $7.12 \times 10^{-2}$ | $7.67 \times 10^{-1}$ | $-4.83 \times 10^{-1}$ | 1.00 |
| Generalised Porous Fisher | 16,000 | $D_0$ | 1.00 | $-4.86 \times 10^{-1}$ | $3.45 \times 10^{-1}$ | $4.60 \times 10^{-1}$ |
| | | $\lambda$ | $-4.86 \times 10^{-1}$ | 1.00 | $-7.65 \times 10^{-1}$ | $2.96 \times 10^{-1}$ |
| | | $K$ | $3.45 \times 10^{-1}$ | $-7.65 \times 10^{-1}$ | 1.00 | $-4.10 \times 10^{-1}$ |
| | | $r$ | $4.60 \times 10^{-1}$ | $2.96 \times 10^{-1}$ | $-4.10 \times 10^{-1}$ | 1.00 |
| Generalised Porous Fisher | 20,000 | $D_0$ | 1.00 | $-7.72 \times 10^{-1}$ | $6.35 \times 10^{-1}$ | $9.03 \times 10^{-1}$ |
| | | $\lambda$ | $-7.72 \times 10^{-1}$ | 1.00 | $-9.10 \times 10^{-1}$ | $-5.88 \times 10^{-1}$ |
| | | $K$ | $6.35 \times 10^{-1}$ | $-9.10 \times 10^{-1}$ | 1.00 | $4.15 \times 10^{-1}$ |
| | | $r$ | $9.03 \times 10^{-1}$ | $-5.88 \times 10^{-1}$ | $4.15 \times 10^{-1}$ | 1.00 |

### D.2 Bivariate marginal posterior PDFs

In the main text we computed only univariate marginal posterior PDFs, we extend this analysis by providing bivariate marginal PDFs here. For the Fisher–KPP and Porous Fisher models we have three bivariate marginal posterior PDFs,

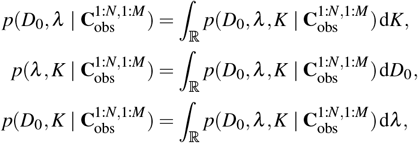

where 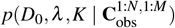 is as given in Equation (17). Similarly, for the Generalised Porous Fisher Model, we have six bivariate marginal posterior PDFs,

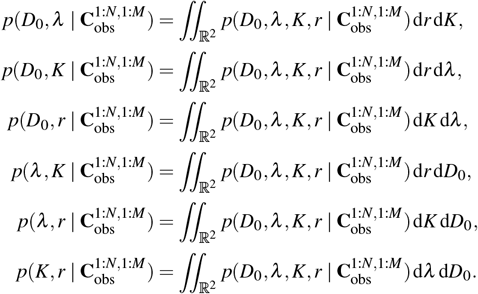

The resulting PDFs using the three initial density conditions are shown for: the Fisher–KPP model (Figure D.2); the Porous Fisher model (Figure D.2); and the Generalised Porous model (Figure D.2).

**Fig. D.1.**
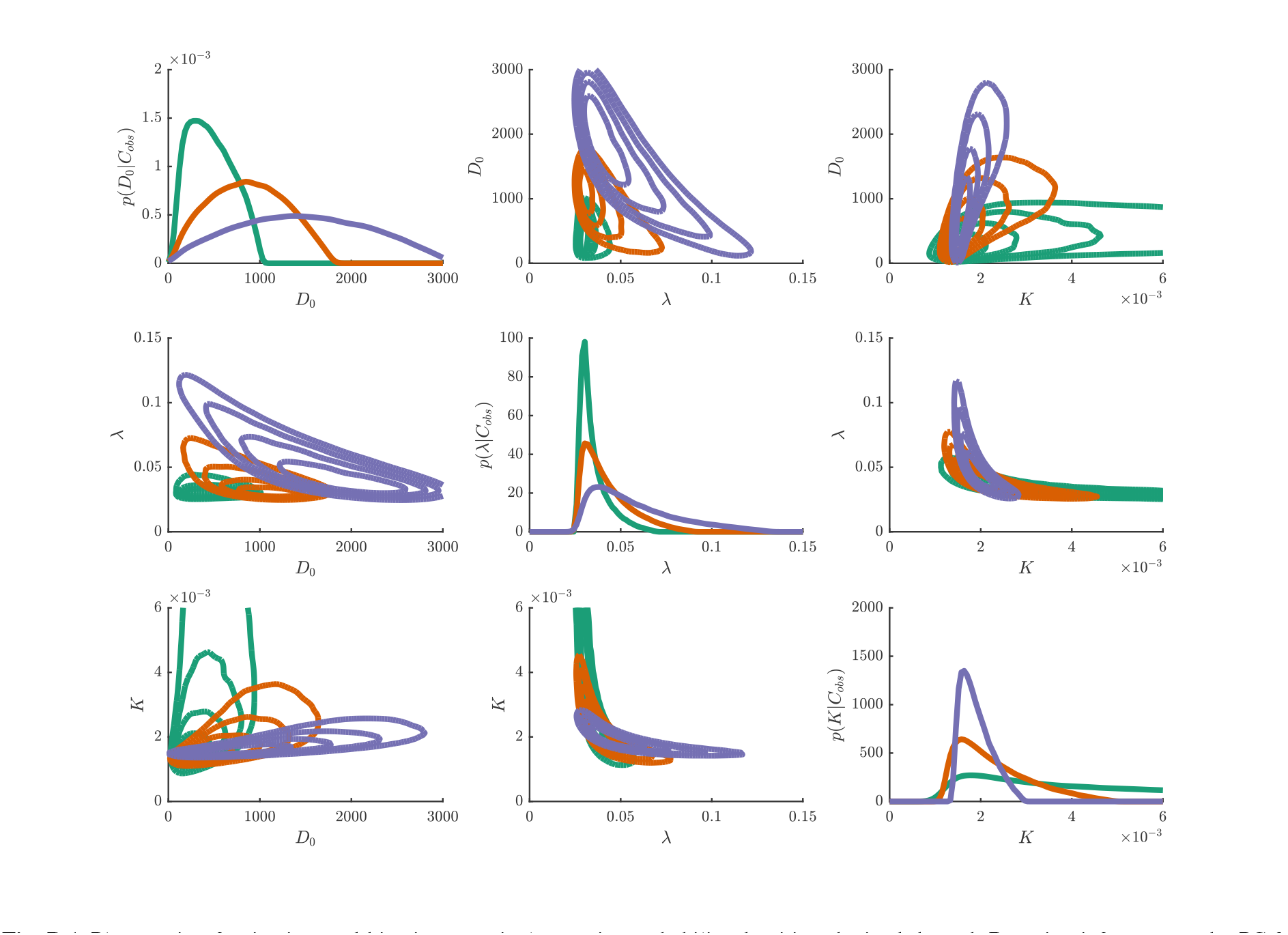
Plot matrix of univariate and bivariate marginal posterior probability densities obtained through Bayesian inference on the PC-3 scratch assay data under the Fisher–KPP model for the three different initial conditions; 12, 000 cells (solid green), 16, 000 cells (solid orange), 20, 000 cells (solid purple). Univariate marginal densities, on the main plot matrix diagonal, demonstrate the degree of uncertainty in the diffusivity, *D*_0_, the proliferation rate, *λ*, and the carrying capacity, *K*. Off diagonals are contour plots of the pairwise bivariate posterior PDFs, these demonstrate the relationships between parameters.

**Fig. D.2.**
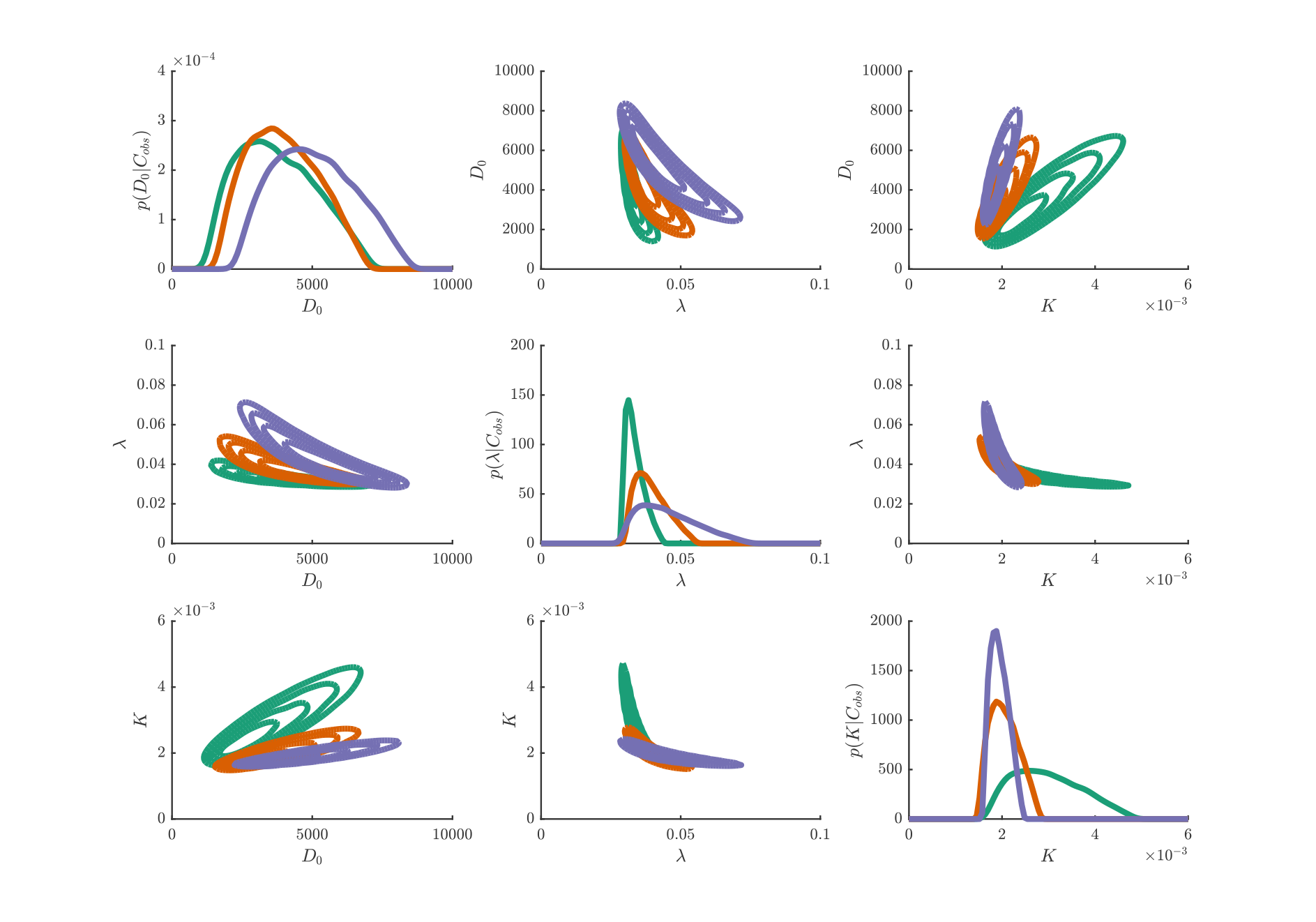
Plot matrix of univariate and bivariate marginal posterior probability densities obtained through Bayesian inference on the PC-3 scratch assay data under the Porous Fisher model for the three different initial conditions; 12, 000 cells (solid green), 16, 000 cells (solid orange), 20, 000 cells (solid purple). Univariate marginal densities, on the main plot matrix diagonal, demonstrate the degree of uncertainty in the diffusivity, *D*_0_, the proliferation rate, *λ*, and the carrying capacity, *K*. Off diagonals are contour plots of the pairwise bivariate posterior PDFs, these demonstrate the relationships between parameters.

**Fig. D.3.**
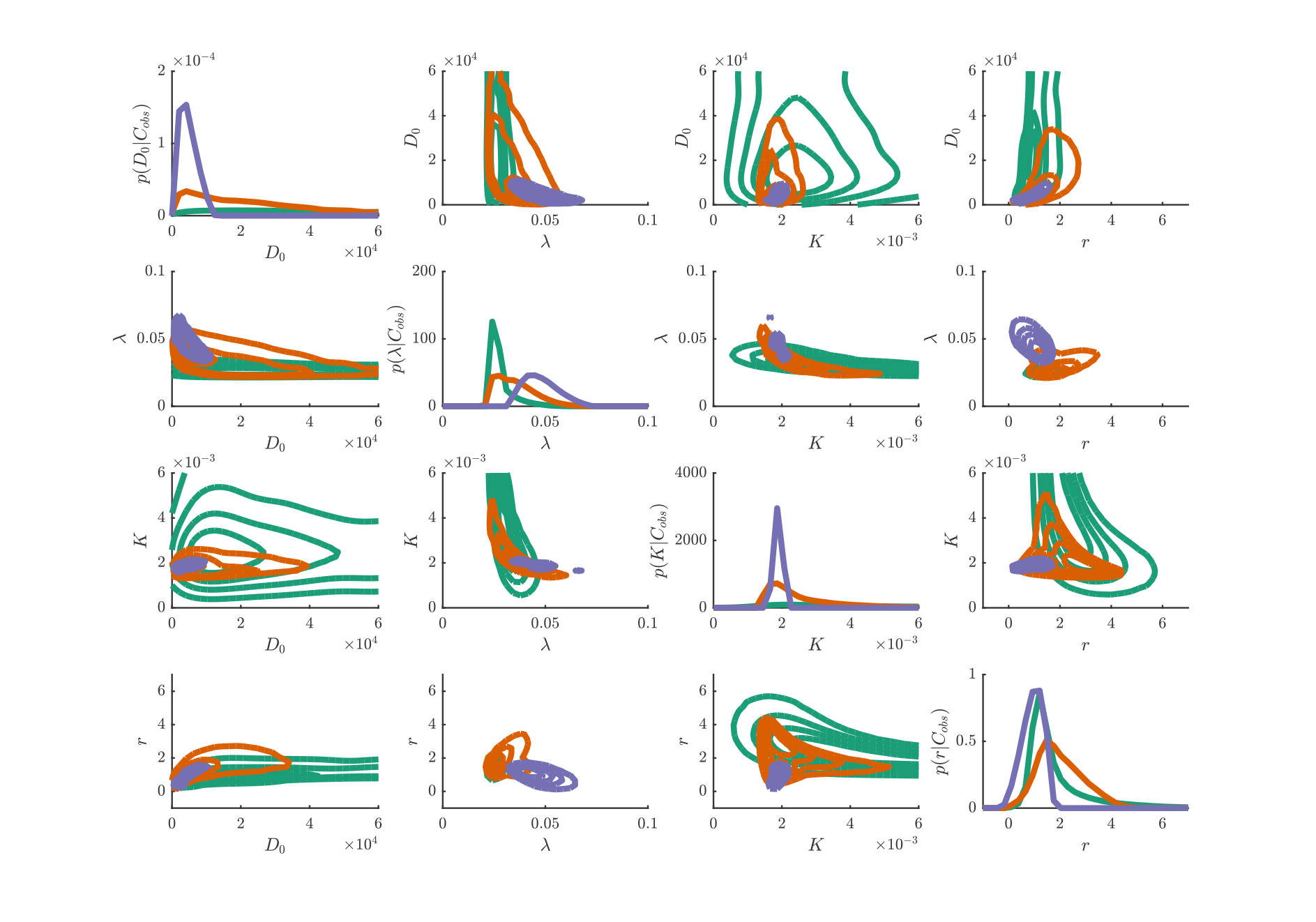
Plot matrix of univariate and bivariate marginal posterior probability densities obtained through Bayesian inference on the PC-3 scratch assay data under the Porous Fisher model for the three different initial conditions; 12, 000 cells (solid green), 16, 000 cells (solid orange), 20, 000 cells (solid purple). Univariate marginal densities, on the main plot matrix diagonal, demonstrate the degree of uncertainty in the diffusivity, *D*_0_, the proliferation rate, *λ*, the carrying capacity, *K*, and the power, *r*. Off diagonals are contour plots of the pairwise bivariate posterior PDFs, these demonstrate the relationships between parameters.

### D.3 Uncertainty in initial condition

In the main text, the assumption was made that *C*_obs_(*x*, 0) = *C*(*x*, 0; ***θ***). That is, we use initial observations as the initial density profile to simulate the model given parameters ***θ***. Since the model is deterministic, the final form of the likelihood is a multivariate Gaussian distribution, which simplifies calculations considerably. Both Jin et al. (2016b) and Warne et al. (2017) indicate that such an assumption could result in underestimation of the uncertainties in parameter estimates.

Following from Warne et al. (2017), we take *C*_obs_(*x*, 0) = *C*(*x*, 0; ***θ***) + *η*_0_, where *η*_0_ is a Gaussian random variable with mean *C*(*x*, 0; ***θ***) and variance 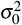. Note that we do not require *σ*_0_ = *σ*, in fact, there are many reasons to consider *σ*_0_ *> σ*. Since 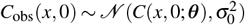, it is also true that 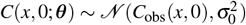. Therefore, our models are to be treated as random PDEs with deterministic dynamics, but random initial conditions.

Since the initial conditions are random, the initial condition is a latent variable that must be integrated out. Thus, the likelihood becomes

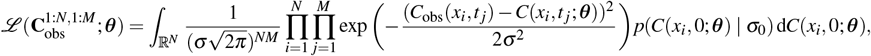

where *σ*_0_ is assumed to be known and *p*(*C*(*x_i_*, 0; ***θ***) *| σ*_0_) is a Gaussian PDF with mean *C*_obs_(*x_i_*, 0) and variance *σ*_0_. This likelihood integral must be computed using Monte Carlo methods. Computationally, we apply directly the ABC MCMC method as given in Algorithm C.2. The only algorithmic difference being that simulated data, *D_s_*, is generated though solving the model PDE after a realisation of the initial density profile has been generated. Overall, this leads to slower convergence in the Markov chain and hence longer computation times.

The inference problem using random initial density profiles was solved using ABC MCMC under the Fisher–KPP model and the Porous Fisher model for initial densities based on 16, 000 initial cells only. We take *σ*_0_ = 2*σ*. Univariate and bivariate marginal posterior PDFs are shown in Figures D.4 and D.5. In the Fisher–KPP model, the additional uncertainty seems to have a significant effect on the uncertainty in the carrying capacity, *K*, in agreement with Warne et al. (2017). However, the diffusion coefficient, *D*_0_, and proliferation rate, *λ*, are not affected as significantly.

**Fig. D.4.**
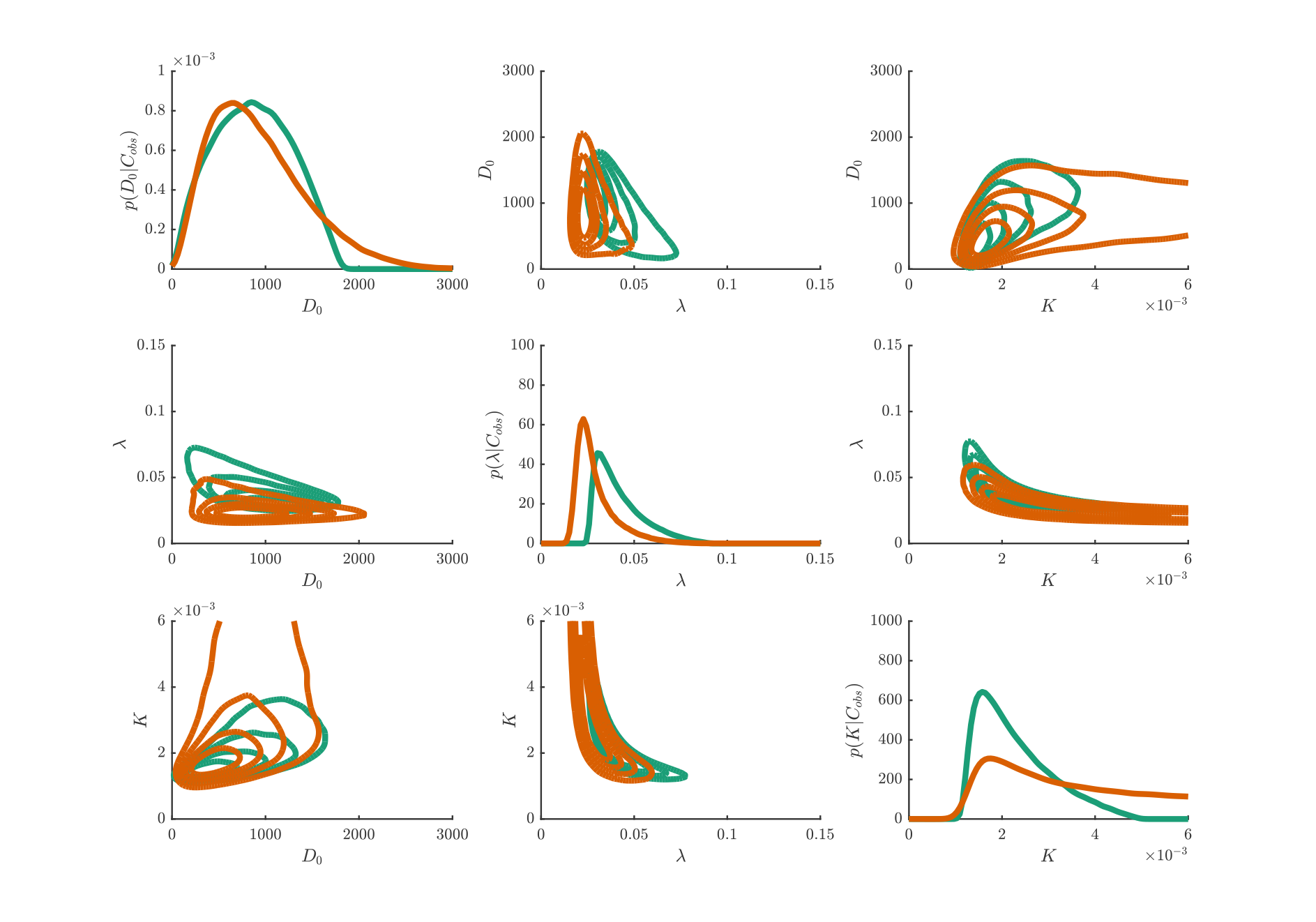
Plot matrix of univariate and bivariate marginal posterior probability densities obtained through Bayesian inference on the PC-3 scratch assay data under the Fisher–KPP model for the fixed initial conditions (solid green) and random initial conditions (solid orange). Univariate marginal densities, on the main plot matrix diagonal, demonstrate the degree of uncertainty in the diffusivity, *D*_0_, the proliferation rate, *λ*, and the carrying capacity, *K*. Off diagonals are contour plots of the pairwise bivariate posterior PDFs, these demonstrate the relationships between parameters.

**Fig. D.5.**
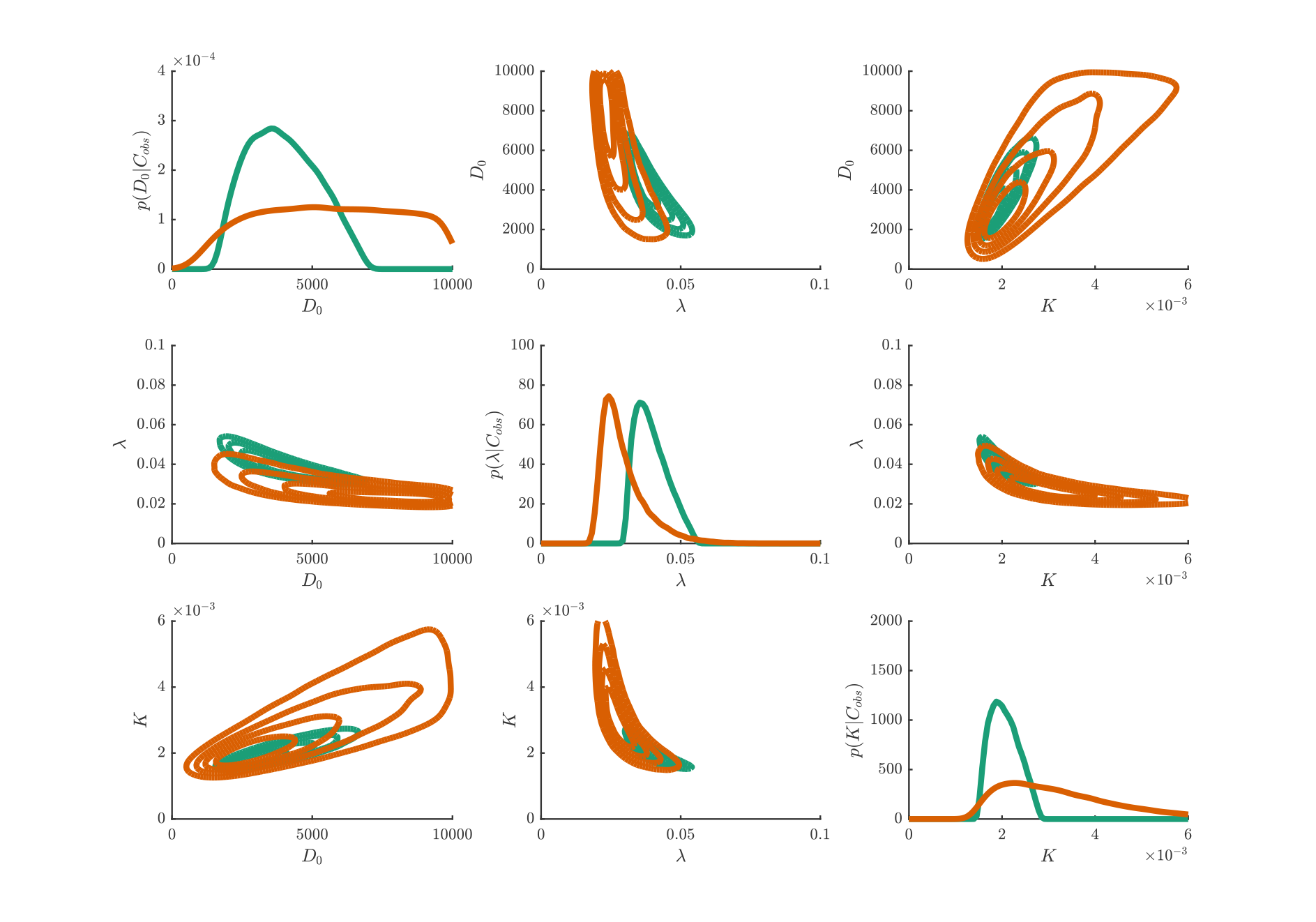
Plot matrix of univariate and bivariate marginal posterior probability densities obtained through Bayesian inference on the PC-3 scratch assay data under the Porous Fisher model for the fixed initial conditions (solid green) and random initial conditions (solid orange). Univariate marginal densities, on the main plot matrix diagonal, demonstrate the degree of uncertainty in the diffusivity, *D*_0_, the proliferation rate, *λ*, and the carrying capacity, *K*. Off diagonals are contour plots of the pairwise bivariate posterior PDFs, these demonstrate the relationships between parameters.

For the Porous Fisher model, both *D*_0_ and *K* are greatly affected. This is not surprising, since motility is density dependent for the Porous Fisher model. By contrast the Fisher–KPP model is almost unaffected in the marginal posterior PDF of *D*_0_, since it is independent of initial cell density.

## Footnotes

1 Many reject the notion that a true model exists (Box, 1976; Spiegelhalter et al., 2014). However, the concept is a useful one for the purposes of deriving information criteria (Akaike, 1974; Spiegelhalter et al., 2002).

## References

Akaike H (1974) A new look at the statistical model identification. IEEE T Automat Contr 19:716–723

Armstrong NJ, Painter KJ, Sherratt JA (2009) Adding adhesion to a chemical signaling model for somite formation. Bull Math Biol 71:1–24

Barenblatt GI (2003) Scaling. Cambridge University Press, Cambridge, UK

Bianchi A, Painter KJ, Sherratt JA (2016) Spatio-temporal models of lymphangiogenesis in wound healing. Bull Math Biol 78:1904–1941

Box GEP (1976) Science and statistics. J Am Stat Assoc 71:791–799

Browning AP, McCue SW, Simpson MJ (2017) A Bayesian computational approach to explore the optimal duration of a cell proliferation assay. Bull Math Biol 79:1888–1906

Cai AQ, Landman KA, Hughes BD (2007) Multi-scale modeling of a wound-healing cell migration assay. J Theor Biol 245:576–594

Crank J (1975) The Mathematics of Diffusion. Oxford University Press, Oxford, UK

Efron B (1986) Why isn’t everyone a Bayesian? Am Stat 40:1–5

Flegg JA, Byrne HM, McElwain DLS (2010) Mathematical model of hyperbaric oxygen therapy applied to chronic diabetic wounds. Bull Math Biol 72:1867–1891

Flegg JA, McElwain DLS, Byrne HM, Turner IW (2009) A three species model to simulate application of hyperbaric oxygen therapy to chronic wounds. PLOS Comput Biol 5:e1000451

Gelman A (2008a) Objections to Bayesian statistics. Bayesian Anal 3:445–450

Gelman A (2008) Rejoinder. Bayesian Anal 3:467–478

Gelman A, Carlin JB, Stern HS, Dunson DB, Vehtari A, Rubin DB (2014) Bayesian Data Analysis, 3rd edn. Chapman & Hall/CRC

Gelman A, Carlin JB, Stern HS, Rubin DB (2004) Bayesian Data Analysis, 2nd edn. Chapman & Hall/CRC

Gerlee P (2013) The model muddle: In search of tumor growth laws. Cancer Res 73:2407–2411

Gurney W, Nisbet R (1975) The regulation of inhomogeneous populations. J Theor Biol 52:441–457

Haridas P, McGovern JA, McElwain DLS, Simpson MJ (2017) Quantitative comparison of the spreading and invasion of radial growth phase and metastatic melanoma cells in a three-dimensional human skin equivalent model. PeerJ 5:e3754

Harris S (2004) Fisher equation with density-dependent diffusion: Special solutions. J Phys A-Math Gen 37:6267

Jackson PR, Juliano J, Hawkins-Daarud A, Rockne RC, Swanson KR (2015) Patient-specific mathematical neurooncology: Using a simple proliferation and invasion tumor model to inform clinical practice. Bull Math Biol 77:846–856

Jin W, Penington CJ, McCue SW, Simpson MJ (2016a) Stochastic simulation tools and continuum models for describing two-dimensional collective cell spreading with universal growth functions. Phys Biol 13:056003

Jin W, Shah ET, Penington CJ, McCue SW, Chopin LK, Simpson MJ (2016b) Reproducibility of scratch assays is affected by the initial degree of confluence: Experiments, modelling and model selection. J Theor Biol 390:136–145

Johnson JB, Omland KS (2004) Model selection in ecology and evolution. Trends Ecol Evol 19:101–108

Johnston ST, Ross JV, Binder BJ, McElwain DLS, Haridas P, Simpson MJ (2016) Quantifying the effect of experimental design choices for in vitro scratch assays. J Theor Biol 400:19–31

Johnston ST, Shah ET, Chopin LK, McElwain DLS, Simpson MJ (2015) Estimating cell diffusivity and cell proliferation rate by interpreting IncuCyte ZOOM™ assay data using the Fisher–Kolmogorov model. BMC Sys Biol 9:38

King JR, McCabe PM (2003) On the Fisher–KPP equation with fast nonlinear diffusion. P Roy Soc Lond A Mat 459:2529–2546

Kullback S, Leibler RA (1951) On information and sufficiency. Ann Math Stat 22:79–86

Lambert B (2018) A Student’s Guide to Bayesian Statistics, 1st edn. Sage Publications

Lambert B, MacLean AL, Fletcher AG, Combes AN, Little MH, Byrne HM (2018) Bayesian inference of agent-based models: A tool for studying kidney branching morphogenesis. J Math Biol 76:1673–1697

Landman KA, Cai AQ, Hughes BD (2007) Travelling waves of attached and detached cells in a wound-healing cell migration assay. Bull Math Biol 69:2119–2138

Liang CC, Park A, Guan JL (2007) In vitro scratch assay: A convenient and inexpensive method for analysis of cell migration in vitro. Nat Protoc 2:329–333

Liepe J, Filippi S, Komorowski M, Stumpf MPH (2013) Maximizing the information content of experiments in systems biology. PLOS Comput Biol 9:e1002888

Maini P, McElwain DS, Leavesley D (2004) Traveling wave model to interpret a wound-healing cell migration assay for human peritoneal mesothelial cells. Tissue Eng 10:475–482

Marchant BP, Norbury J, Sherratt JA (2001) Travelling wave solutions to a haptotaxis-dominated model of malignant invasion. Nonlinearity 14:1653–1671

Marjoram P, Molitor J, Plagnol V, Tavaré S (2003) Markov chain Monte Carlo without likelihoods. Proc Natl Acad Sci USA 100:15324–15328

Matsiaka OM, Baker RE, Shah ET, Simpson, MJ (2018) Mechanistic and experimental models of cell migration reveal the importance of intercellular interactions in cell invasion. bioRxiv preprint doi:10.1101/391557

Murray JD (2002) Mathematical Biology: I. An Introduction. Springer, New York

Nardini JT, Chapnick DA, Liu X, Bortz DM (2016) Modeling keratinocyte wound healing dynamics: Cell-cell adhesion promotes sustained collective migration. J Theor Biol 400:103–117

Parker A, Simpson MJ, Baker RE. The impact of experimental design choices on parameter inference for models of growing cell colonies. Roy Soc Open Sci 5:180384

Pooley CM, Marion G (2018) Bayesian model evidence as a practical alternative to deviance information criterion. Roy Soc Open Sci 5:171519

Sarapata EA, de Pillis LG (2014) A comparison and catalog of intrinsic tumor growth models. Bull Math Biol 76:2010–2024

Savla U, Olson LE, Waters CM (2004) Mathematical modeling of airway epithelial wound closure during cyclic mechanical strain. J Appl Physiol 96:566–574

Schwarz G (1978) Estimating the dimension of a model. Ann Stat 6:461–464

Sengers BG, Please CP, Oreffo RO (2007) Experimental characterization and computational modelling of two– dimensional cell spreading for skeletal regeneration. J R Soc Interface 4:1107–1117

Shannon CE (1948) A mathematical theory of communication. Bell Syst Tech J 27:379–423

Sherratt JA, Murray JD (1990) Models of epidermal wound healing. P Roy Soc Lond B Bio 241:29–36

Silk D, Kirk PDW, Barnes CP, Toni T, Stumpf MPH (2014) Model selection in systems biology depends on experimental design. PLOS Comput Biol 10:e1003650

Silverman BW (1986) Density Estimation for Statistics and Data Analysis. Chapman & Hall/CRC

Simpson MJ, Baker RE, McCue SW (2011) Models of collective cell spreading with variable cell aspect ratio: A motivation for degenerate diffusion models. Phys Rev E 83:021901

Simpson MJ, Landman KA, Hughes BD, Newgreen DF (2006) Looking inside an invasion wave of cells using continuum models: Proliferation is the key. J Theor Biol 243:343–360

Simpson MJ, Zhang DC, Mariani M, Landman KA, Newgreen DF (2007) Cell proliferation drives neural crest cell invasion of the intestine. Dev Biol 302:553–568

Sisson SA, Fan Y, Tanaka MM (2007) Sequential Monte Carlo without likelihoods. Proc Natl Acad Sci USA 104:1760–1765

Slezak F, Diego, Surez C, Cecchi GA, Marshall G, Stolovitzky G (2010) When the optimal is not the best: Parameter estimation in complex biological models. PLOS ONE 5:e13283

Spiegelhalter DJ, Best NG, Carlin BP, Van Der Linde A (2002) Bayesian measures of model complexity and fit. J Roy Stat Soc B 64:583–639

Spiegelhalter DJ, Best NG, Carlin BP, van der Linde A (2014) The deviance information criterion: 12 years on. J Roy Stat Soc B 76:485–493

Stoica P, Selen Y (2004) Model-order selection: A review of information criterion rules. IEEE Signal Proc Mag 21:36–47

Sunnåker M, Busetto AG, Numminen E, Corander J, Foll M, Dessimoz C (2013) Approximate Bayesian computation. PLOS Comput Biol 9:e1002803

Swanson KR, Alvord EC, Murray JD (2002) Virtual brain tumours (gliomas) enhance the reality of medical imaging and highlight inadequacies of current therapy. Br J Cancer 86:14–18.

Swanson KR, Bridge C, Murray JD, Alvord EC (2003) Virtual and real brain tumors: Using mathematical modeling to quantify glioma growth and invasion. J Neurol Sci 216:1–10

Tsoularis A, Wallace J (2002) Analysis of logistic growth models. Math Biosci 179:21–55

Vanlier J, Tiemann CA, Hilbers PAJ, van Riel NAW (2012) A Bayesian approach to targeted experiment design. Bioinformatics 28:1136–1142

Vittadello ST, McCue SW, Gunasingh G, Haass NK, Simpson MJ (2018) Mathematical models for cell migration with real-time cell cycle dynamics. Biophys J 114:1241–1253

Warne DJ, Baker RE, Simpson MJ (2017) Optimal quantification of contact inhibition in cell populations. Biophys J 113:1920–1924

Warne DJ, Baker RE, Simpson MJ (2018) Multilevel rejection sampling for approximate Bayesian computation. Comput Stat Data An 124:71–86

Wilkinson RD (2013) Approximate Bayesian computation (ABC) gives exact results under the assumption of model error. Stat Appl Genet Mol 12:129–141

Witelski TP (1995) Merging traveling waves for the porous-Fisher’s equation. Appl Math Lett 8:57–62

Yang Y (2005) Can the strengths of AIC and BIC be shared? a conflict between model identification and regression estimation. Biometrika 92:937–950

